# Characterization of the subunit composition and structure of native adult glycine receptors

**DOI:** 10.1101/2021.05.17.444520

**Authors:** Hailong Yu, Xiao-chen Bai, Weiwei Wang

**Affiliations:** Department of Biophysics, University of Texas Southwestern Medical Center, Dallas, TX 75390, USA

**Keywords:** heteromeric glycine receptor, Cryo-EM structure, stoichiometry, planar bilayer recording, stepwise photobleaching

## Abstract

The strychnine-sensitive pentameric Glycine Receptor (GlyR) mediates fast inhibitory neurotransmission in the mammalian nervous system. Only heteromeric GlyRs mediate synaptic transmission, since they contain the β subunit that permits clustering at the synapse through its interaction with scaffolding proteins. Here we show that α2 and β subunits assemble with an unexpected 4:1 stoichiometry to produce GlyR with native electrophysiological properties. We determined structures in multiple functional states at 3.6 – 3.8 Å resolutions and show α2β GlyR assembly mechanism. Furthermore, we show that one single β subunit in each GlyR gives rise to the characteristic electrophysiological properties of heteromeric GlyR, while more β subunits renders GlyR non-conductive. A single β subunit ensures a univalent GlyR-scaffold linkage, which means the scaffold alone regulates the cluster properties.

## INTRODUCTION

Nervous system function depends on electrical signal transmission across neuronal cells. At chemical synapses, electrical signals are transmitted through the release of neurotransmitters from the pre-synaptic neuron and resultant activation of the cognate neurotransmitter receptors in the post-synaptic neuron (Eccles, 1982). The neurotransmitter glycine mediates fast inhibitory neurotransmission through activating the strychnine-sensitive glycine receptors (GlyRs) at inhibitory synapses (Moss and Smart, 2001; Pfeiffer and Betz, 1981; Werman et al., 1967). GlyRs are expressed in both the central and peripheral nervous systems(Lynch, 2004). Defects in GlyRs cause the neurological disorder hyperekplexia (Bode and Lynch, 2014) and are implicated in autism spectrum disorders (Yu et al., 2013). GlyRs have also been found in the visual system (Wassle et al., 2009) and identified as possible therapeutic targets for inflammatory pain (Harvey et al., 2004; Lynch and Callister, 2006).

GlyRs are members in the ionotropic pentameric Cys-loop receptor superfamily. There are two major types of GlyRs, homomeric GlyRs composed of only α subunits and heteromeric GlyRs consisting of both α and β subunits. Extensive study on the homomeric GlyRs (Du et al., 2015; Huang et al., 2015; Kumar et al., 2020; Yu et al., 2021) and related Cys-loop receptors (Phulera et al., 2018; Rahman et al., 2020; Zhu et al., 2018) provided little information on the architecture and working mechanism of native adult GlyRs for the following reasons. First, homomeric GlyRs are found almost exclusively during early development while heteromeric GlyRs is the predominantly form in adult animals (Malosio et al., 1991; Weltzien et al., 2012). Second, heteromeric GlyRs exhibit very different electrophysiological properties, which is solely determined by the β subunit that occur only in the heteromeric GlyRs (Bormann et al., 1993; Lynch, 2009; Pribilla et al., 1992). Third, heteromeric GlyRs form high-density clusters at post-synaptic membrane and mediate synaptic transmission while homomeric GlyRs do not localize at synapses (Kirsch et al., 1993; Specht et al., 2013). This is because homomeric GlyRs do not contain the β subunit with which post-synaptic scaffolding protein gephyrin directly associates (Meyer et al., 1995).

We know very little about heteromeric GlyRs that even the subunit composition has been controversial for decades (Durisic et al., 2012; Grudzinska et al., 2005; Kuhse et al., 1993; Langosch et al., 1988; Patrizio et al., 2017; Yang et al., 2012). The number and spatial arrangement of β subunits in each GlyRs are central questions in synaptic transmission since they impact how heteromeric GlyR clusters with multivalent scaffolding protein gephyrin (Sander et al., 2013; Sola et al., 2004) and regulate synaptic plasticity (Citri and Malenka, 2008; Zacchi et al., 2014; Zeng et al., 2018). Here, we characterized the function of heteromeric α2β GlyR both in cells and in total reconstitution systems. A combination of biochemical and structural analyses revealed the molecular mechanism underlying the assembly of α2β GlyR consisting four α2 and one β subunit, as well as how such molecular composition gives rise to native GlyR electrophysiological characteristics, including glycine activation, single-channel conductance and picrotoxin (PTX) inhibition. We further show how α2β GlyR containing more than one β subunit does not conduct Cl^-^ upon glycine activation. Our findings lead to the conclusion that a 4:1 α2:β stoichiometry accounts for the native physiology of heteromeric GlyRs.

## RESULTS

### Invariant α2β GlyR subunit stoichiometry

We produced heteromeric α2β GlyR through co-expressing the α2 and the β subunits in HEK293 cells for structural and functional characterization. We have identified α2β GlyR constructs, α2em and βem (see methods for details, denoted here α2 and β unless otherwise indicated), that exhibited both good physiological function (Figure S1, A and B) and biochemical behavior, allowing unambiguous characterization of the molecular composition. Depending on the expression levels of each subunit, homomeric α2 GlyR, heteromeric α2β GlyR, or a mixture of both may be produced. We used whole-cell patch-clamp electrophysiology to characterize the GlyRs residing in the plasma membrane as a function of expression levels.

Homomeric GlyRs are inhibited by the convulsant alkaloid, pore-blocking toxin picrotoxin (PTX) in the micro-molar range while native heteromeric GlyRs are more resistant with an over 10-fold higher IC_50_ (Pribilla et al., 1992; Shan et al., 2001). When transfected with only α2, homomeric GlyRs exhibited high affinity to PTX with an IC_50_ of ∼2.4 µM (Figure 1A and 1C black, Figure S1D) (Pribilla et al., 1992; Yang et al., 2007). When a moderate amount of the β subunit was co-expressed, at 1:1 α2:β ratio (transfected DNA amounts, Figure S1C), the IC_50_ increased to ∼ 11 µM (Figure 1C green), indicating that a mixture of α2 and α2β GlyR were produced. When an excess of the β subunit was expressed at 1:3 α2:β ratio, PTX IC_50_ increased further to ∼30 µM (Figure 1, B and C blue). However, a greater excess of the β subunit (at 1:10 α2:β ratio) did not result in further change (Figure 1C red).

**Figure 1.**
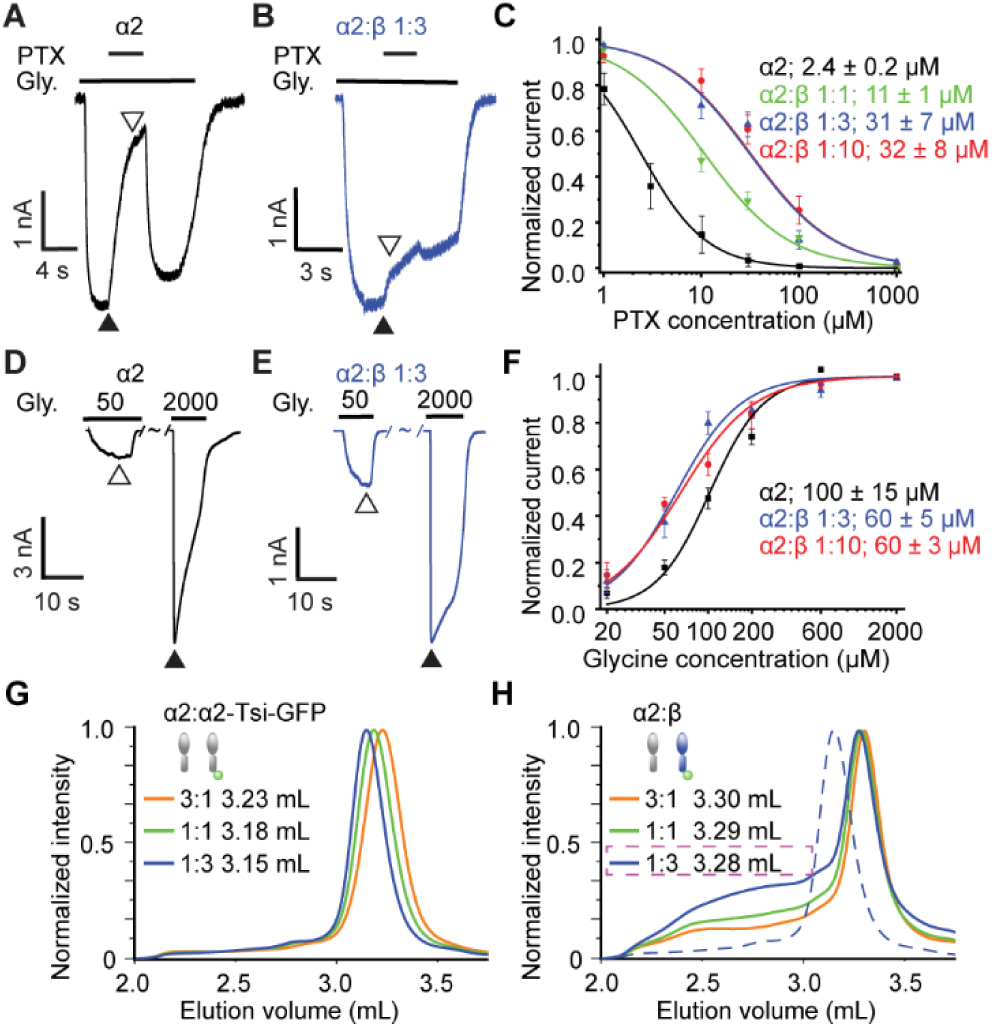
**Saturable effects of β subunit expression levels** (A and B) Representative recordings of GlyR inhibition by 10 μM PTX in the presence of 100 μM glycine at α2:β expression ratios of 1:0 (A) and 1:3 (B). Normalized current was calculated by dividing the currents immediately after (open triangle) PTX application by that before (closed triangle). (C) Dose response of PTX with IC50 resulting from Hill fits (lines) listed. Data are represented as mean ± SEM (n = 3-6 cells). (D and E) Glycine activation at α2:β ratios of 1:0 (D) and 1:3 (E).Glycine concentrations are in μM. (F) Dose response of glycine with EC50 resulting from Hill fits (lines) listed. Data are represented as mean ± SEM (n = 3-6 cells). (G and H) FSEC elution profiles of (G) α2:α2-Tsi-GFP ratio of 3:1 (red), 1:1 (green) and 1:3 (blue) and (H) α2:β ratio of 3:1 (red), 1:1 (green) and 1:3 (blue). Dashed line denotes 1:3 α2:α2- Tsi-GFP.Peak elution volumes are listed. See also Figure S1.

Unlike PTX, neurotransmitter glycine binds to the GlyR extracellular domain, distant from the ion conduction pore. Based on the difference in its affinity for homomeric and heteromeric GlyR (Grudzinska et al., 2005; Pribilla et al., 1992), we used glycine as an additional probe of the formation of α2β GlyR (Figure S1E). When only α2 GlyR was expressed, a ∼100 µM EC_50_ (Figure 1, D and F black) was observed. With both the 1:3 and 1:10 α2:β ratio, the EC_50_ decreased to ∼60 µM (Figure 1, E and F blue, red). These numbers are within the range of reported glycine affinities for α2 and α2β GlyRs (Grudzinska et al., 2005; Mohammadi et al., 2003; Pribilla et al., 1992). Clearly, the change in glycine sensitivity followed a similar trend as that of PTX – the effects of β subunit expression saturate at 1:3 α2:β ratio. This trend has two implications: first, at a 1:3 α2:β ratio, sufficient β subunit is expressed to make α2β GlyR the predominant form. Second, the number of β subunits in each α2β GlyR did not further increase when the α2:β ratio was changed from 1:3 to 1:10, since the incorporation of additional β subunits should further change the PTX and glycine affinities (Pribilla et al., 1992; Shan et al., 2001).

We further used Fluorescence Size Exclusion Chromatography (FSEC) to test whether the number of β subunit in each α2β GlyR stayed the same irrespective of expression levels of α2 and β subunits. Since the β subunit has a higher molecular weight than that of α2, changes in α2:β stoichiometry should result in changes of elution volumes. As control experiments, we co-expressed α2, and the higher molecular weight fluorescent fusion α2-Tsi-GFP (see methods), over a wide range of expression ratios (virus MOI, see methods) (Morales-Perez et al., 2016). A clear shift in the peak elution volume was observed (Figure 1G and Figure S1F, left). This finding supports that α subunits are able to form GlyRs with arbitrary stoichiometries (Griffon et al., 1999; Kuhse et al., 1993). In contrast, when co-expressing α2 and β, the peak elution volume did not change over the same range of virus ratios (Figure 1H and Figure S1F, right). These observations point to an invariant subunit stoichiometry irrespective of expression levels of α2 and β, consistent with our and previously reported electrophysiology data (Durisic et al., 2012; Kuhse et al., 1993). Clearly, GlyRs are unlike other members of the Cys-loop receptor family in which variability in stoichiometry has been shown (Morales-Perez et al., 2016).

### Planar lipid bilayer electrophysiology

Although electrophysiological properties of recombinantly expressed homomeric GlyR have been characterized (Cascio et al., 2001), functional reconstitution of heteromeric GlyR has not been reported. We employed the planar lipid bilayer method to characterize the electrophysiology of purified α2β GlyR *in vitro* (Figure 2A) (Mueller et al., 1962; Wang et al., 2014). In this method, a synthetic bilayer was created using the painting method across a ∼100 µm hole. Purified α2 (α2-Tsi-GFP) and α2β (expressed with 1:3 α2:β ratio) GlyRs were reconstituted into proteoliposomes and subsequently incorporated into the synthetic membrane through salt-assisted spontaneous fusion. GlyR electrophysiological properties were thus characterized in a chemically defined environment.

**Figure. 2.**
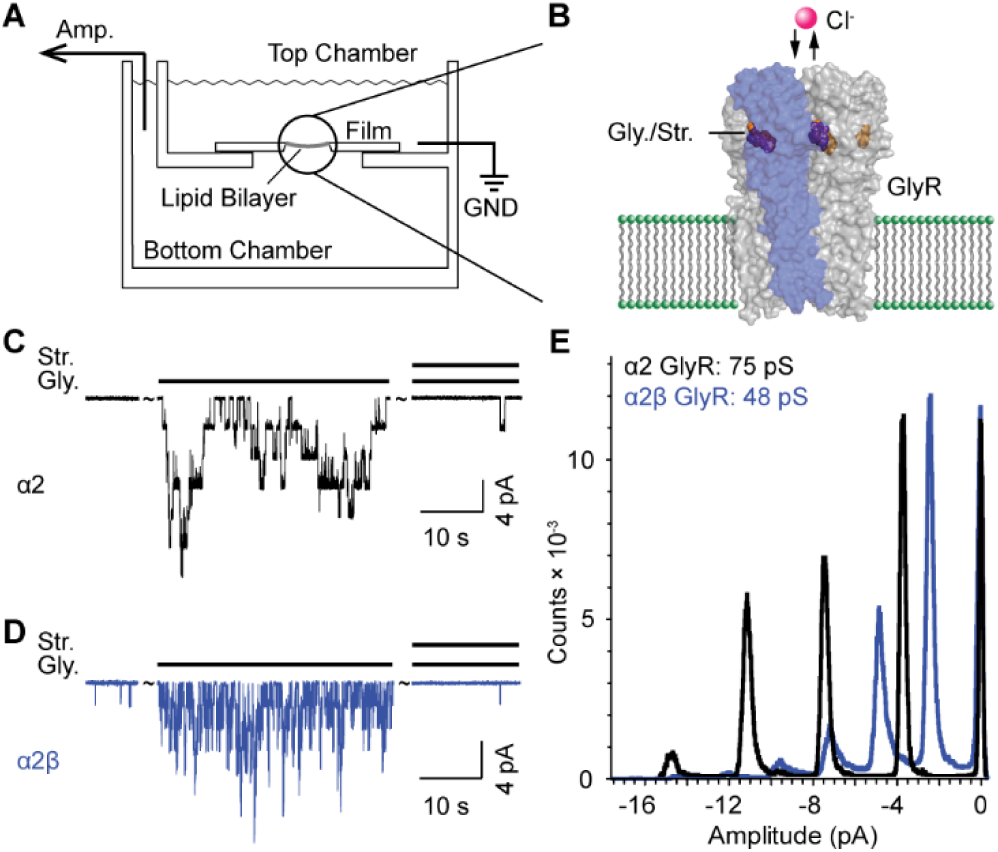
**Electrophysiology of GlyRs in planar lipid bilayers.** (A and B) Illustration of (A) the recording apparatus and (B) GlyR reconstituted into artificial bilayers. (C and D) Current traces of (C) α2 and (D) α2β GlyRs at -50 mV membrane potential. Solution on both sides contained (in mM): 10 potassium phosphate pH 7.4, 150 KCl and 1 EDTA. Glycine (Gly.) and strychinine (Str.) concentrations were 200 µM and 10 µM, respectively. (E) Histogram of all points from traces of α2 homomer (color in black) and α2β heteromer (color in blue). The conductance of α2 and α2β heteromer are listed.

The α2 GlyR exhibited rare spontaneous openings in the absence of ligands (Figure 2C left). When 200 µM glycine was applied to the same membrane, robust channel activation was observed (Figure 2C middle). This observation is consistent with the low ‘basal’ activity and strong glycine activation found in cellular membranes (Schmieden et al., 1989; Sontheimer et al., 1989) (See Figure 1). A histogram of the current levels (Figure 2E black) led to the identification of a ∼75 pS single-channel conductance. This value is consistent with one major conductance observed in cells (∼ 70 pS) (Bormann et al., 1993; Cascio et al., 1993; Lynch, 2009). Subsequent application of 10 µM competitive antagonist strychnine(Becker et al., 1988) on the same side of the membrane almost completely inhibited α2 GlyR activity (Figure 2C right). Very similar pharmacology – low basal activity (Figure 2D left), strong glycine activation (Figure 2D middle) and strychnine inhibition (Figure 2D right) – was observed for the α2β GlyR. However, the α2β GlyR had a much lower single-channel conductance of ∼48 pS (Figure 2E blue), which recapitulates the major conductance of heteromeric GlyRs both heterologously expressed and in native adult tissues (∼50 pS) (Ali et al., 2000; Bormann et al., 1993; Lynch, 2009; Takahashi et al., 1992). The single channel conductance of both α2 and α2β GlyRs remained constant in bilayers made with either polar lipids extracted from brain tissues or an artificial mimic (see Methods), indicating minimal lipid effects in these mixtures. The uniformity of single channel conductance (Figure 2E) further confirms a homogeneous subunit stoichiometry of the purified α2β GlyR, since the unique sequence features in the M2 helix of the β subunit has been shown to alter GlyR conductance (Bormann et al., 1993). Therefore, based on the similarity of its electrophysiological properties to native heteromeric GlyRs(Ali et al., 2000; Bormann et al., 1993; Lynch, 2009; Takahashi et al., 1992), we believe this purified α2β GlyR is likely to be the dominant, physiological form of α2β GlyR.

### 4:1 α2:β GlyR structures in the closed, desensitized and open states

The α2β GlyR structures were resolved in complex with the neurotransmitter glycine, as well as a competitive antagonist strychnine in saposin nanodiscs (Flayhan et al., 2018). Reconstructed cryo-EM maps contained densities of α2β GlyR (Figure 3A grey), as well as a single extra satellite near the intracellular side (Figure 3A green) resulting from the GFP fusion in the intracellular M3-M4 loop of the β subunit (see methods for details), suggesting that each GlyR contains only one β subunit. Within a mask excluding this satellite density (Figure 3A translucent blue surface), the α2β GlyR maps reached overall resolutions of 3.6 – 3.8 Å (Figure S2), allowing for unambiguous amino acid registration for the majority of the receptor (Figure S3). One β subunit (Figure 3B blue) and four α2 subunits (Figure 3B grey) were identified in each pentameric α2β GlyR in the atomic models, resulting in an unexpected but unequivocal 4:1 subunit stoichiometry.

**Figure 3.**
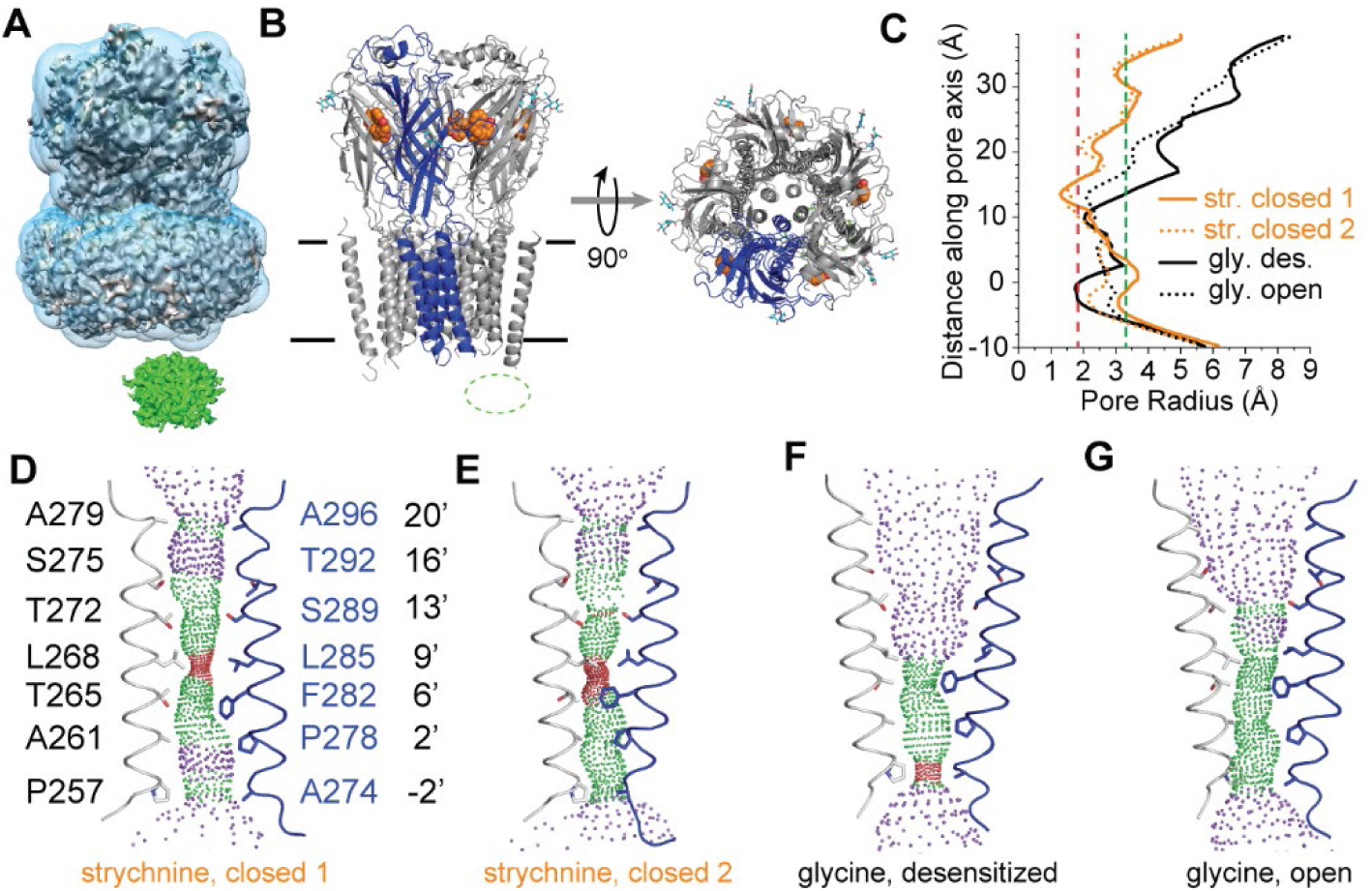
**Overall structures of α2β GlyR in complex with glycine and strychnine.** (A) One α2β GlyR-strychnine cryo-EM density map containing GlyR (grey) and a GFP blob (green), with a mask excluding GFP (translucent blue). (B) Cartoon representations of α2β GlyR atomic model viewed parallel to the membrane (black lines) plane (left) and down the extracellular side (right). Grey: α2, blue: β, orange: strychnine, cyan sticks: N-linked glycans. (C) Pore radii plotted against the distance along the pore axis from the Cα position of 0’ Arginine. (D-G) Ion permeation pathways of (D) strychnine-bound closed 1, (E) closed 2, and (F) glycine-bound desensitized, (G) open states. Purple, green and red mesh represents radii of >3.3 Å, 1.8-3.3 Å and < 1.8 Å, respectively. Sidechains of pore-lining residues are highlighted as sticks. See also Figure S2, S3, S4 and S5.

Two structural classes were present in the GlyR-strychnine complex (Figure S2A). In both classes, the ion-conduction pores were tightly constricted near the 9’ amino acid residues α2: L268 and β: L285 (Figure 3, D and E), too narrow (< 1.8 Å radii) for even non-hydrated Cl^-^ to pass through (Figure 3C orange). The antagonist strychnine was present in all 5 neurotransmitter binding pockets in both structures (Figure S4, A and B left). These binding pockets are in an apo/resting conformation where the loop Cs are in the “uncapped” configuration (Yu et al., 2021) (Figure S5, G and I blue). The differences between these two structures are manifested in the transmembrane domain (TMD) near the intracellular leaflet. One class (closed 1) is pseudo-5-fold symmetrical across the whole receptor (Figure S4, B and C), while the other class (closed 2) has the β TM2 tilted toward the ion conduction pore and one of the α2 TM2 tilted out (Figure S4D). Since both structures exhibit functional characteristics of a closed conduction pore, we believe they represents two possible sub-conformations in the closed state of heteromeric GlyR.

In the α2β GlyR-glycine complex, we also identified two distinct structural classes (Figure S2H). In both classes, the conduction pores near the 9’ position widened and no longer cause constriction as in the GlyR-strychnine structures. The radii were larger than 1.8 Å (Figure 3C black) that allow partially hydrated Cl^-^ to pass through. In one class, the ion conduction pore narrowed near the -2’ amino acid residues α2: P257 and β: A274, resulting in a new constriction near the intracellular side (Figure 3F). Glycine densities were observed in all the 5 neurotransmitter binding pockets (Figure S4B middle) that are in the agonist-bound conformation (Figure S5, D-F). These features suggest that this structural class represents the desensitized functional state. In contrast, an asymmetrical widening near -2’ position in the other structural class removed constriction (Figure 3G and Figure S4F). The resultant pore radii were larger than 1.8 Å all across (Figure 3C). All 5 neurotransmitter binding pockets were in the agonist-bound conformation even though clear densities of glycine were observed for only 3 (Figure S4B right). Since there is no constriction near the 9’ or the -2’ positions and the pore size allows for partially hydrated Cl^-^ to pass through, similar to homomeric GlyRs in the open state (Kumar et al., 2020; Yu et al., 2021), we believe this class represents a structure of α2β GlyR in the open functional state.

### Structural determinants of α2β GlyR assembly and electrophysiology

The α2β GlyR shares similar overall structures with homomeric GlyRs (Du et al., 2015; Kumar et al., 2020; Yu et al., 2021). The constriction sites in the ion conduction pathway, 9’ in the closed state and -2’ in the desensitized state, are similar (Figure 3). In addition, the neurotransmitter binding sites are highly conserved across the α/β, β/α and α/α interfaces and exhibit similar structural changes upon agonist binding (Figure S5). These conservation in binding sites explain the small difference in glycine affinity between α2β and α2 GlyRs (Figure 1F) (Grudzinska et al., 2005; Pribilla et al., 1992). Despite apparent similarities, several features in the β subunit give rise to the unique properties of native GlyRs.

Glycosylation sites of the β subunit are required for proper assembly of α2β GlyR. Based on sequence homology, two N-glycosylation sites have been predicted (Griffon et al., 1999; Schaefer et al., 2018). β:N220 is specific to the β subunit and located to the N-terminus of loop C, closer to the neurotransmitter binding pocket compared with α2 (Figure 4A, Figure S3). The β:N220A mutation abolished glycosylation and prevented proper assembly of the α2β GlyR (Figure S5J). β:N36 was believed homologous to α2:N45 (Griffon et al., 1999; Schaefer et al., 2018) but turned out to locate near the vestibule (Figure 4A and B). Although the resolution was not sufficient for modeling, density maps show that the glycan extends into the vestibule (Figure 4B). β:N36A mutation also prevented expression of the α2β GlyR (Figure S5J), suggesting its essential role in heteromeric GlyR assembly. Glycosylation at this site is likely a reason for the invariant 4:1 α2:β stoichiometry since the vestibule is not large enough to accommodate more than one β:N36 glycan.

**Figure 4.**
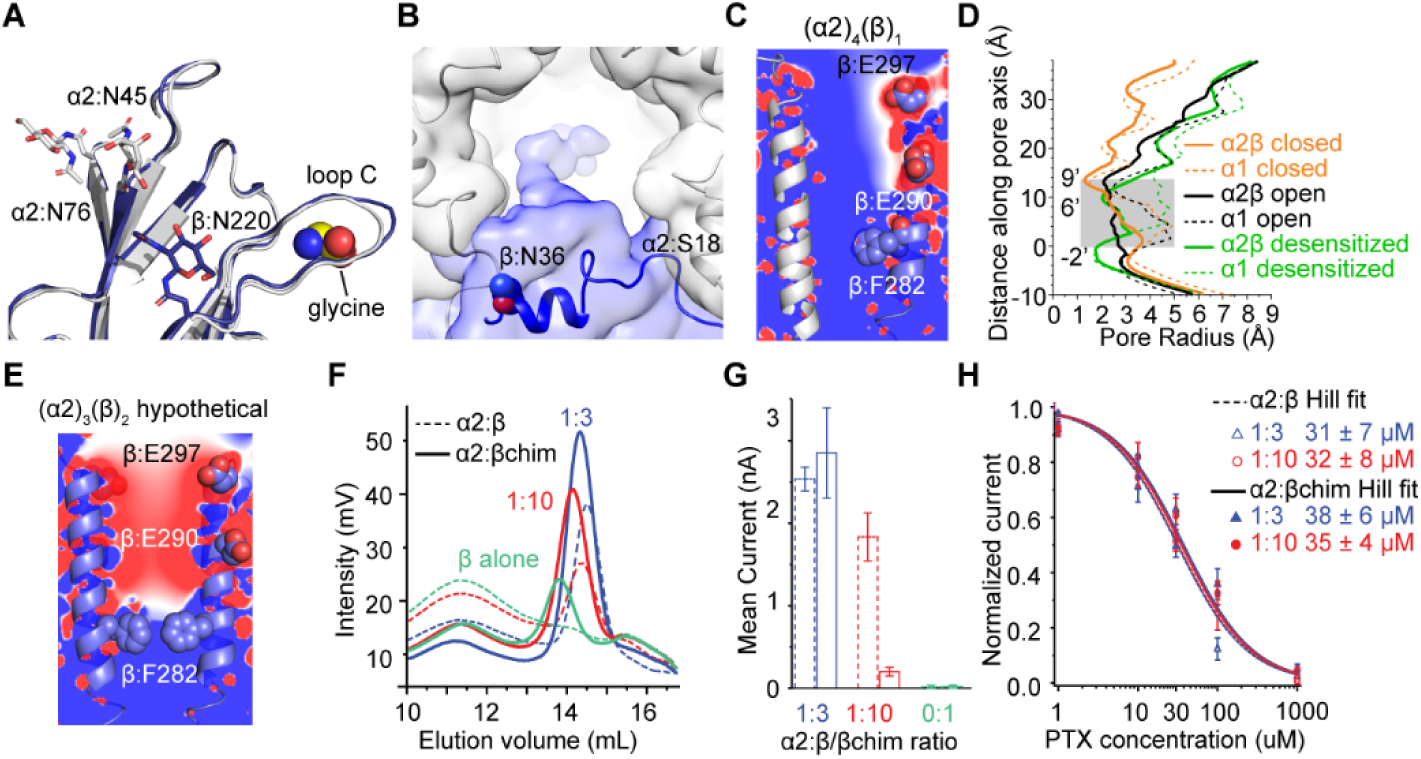
**Features of the β subunit underlying α2β GlyR physiology.** (A) N-linked glycosylation sites on α2 (grey) and β (blue) subunits facing away the vestibule. (B) cryo-EM density originating from β:N36 (blue) and extending into the vestibule. Map is low-pass filtered and 6 Å and shown as translucent surface. (C) Ion permeation pore of α2β GlyR with β: E297, E290 and F282 shown as spheres. Electrical potentials are shown as color gradient from red (-2 kT/e) to blue (2 kT/e). (D) Pore radii in the closed, open and desensitized states of α2β (see Figure 3C) and homomeric α1 GlyRs (PDB ID: 3JAD, 6PM6, 6PM5, respectively). Regions exhibiting clear differences between α2β and α1 are shaded. (E) Ion permeation pore of hypothetical (α2)3(β)2 represented as in (C). (F) FSEC of α2 co-expressed with β (dashed) and βchim (solid) at ratios of 1:3 (blue), 1:10 (red) and β alone (light green). (G) Whole-cell current activated by 100 µM glycine (n = 5-6 cells, mean ± SEM). (H) PTX dose response curves of α2:β and α2:βchim expressed at 1:3 and 1:10 ratios (n = 3-6 cells, mean ± SEM). See also Figure S3, S5 and S6.

The unique amino-acid composition of the pore-lining TM2 of β subunit (Figure S6) resulted in electrophysiological properties specific to heteromeric α2β GlyR. First, two negatively-charged amino-acid residues, β:E297 and β:E290, poised near the extracellular side of transmembrane pore. The resultant negative electrostatic potential (Figure 4C) most likely reduced the local concentration of Cl^-^, decreasing ion conductance. Indeed, the lower single-channel conductance of heteromeric GlyRs (Figure 2E) compared with homomeric GlyRs has been functionally attributed to these residues (Bormann et al., 1993). Second, the bulky side chain of 6’ β:F282 narrows the pore radii in all functional states compared with homomeric GlyRs (Figure 4, C and D). This explains why heteromeric GlyRs exhibit much lower affinity to the pore blocking PTX: PTX binds near 6’ position (Kumar et al., 2020) and β:F282 side chain constitutes steric hindrance. Supporting this conclusion, mutating β:F282 to amino acids with smaller side-chains rendered high-affinity blocking of heteromeric GlyRs just like with homomeric GlyRs (Pribilla et al., 1992).

### Only 4:1 α2:β GlyR conducts

The negative charges of β:E297 and β:E290 and reduced pore radii due to the bulky β:F282 predict that heteromeric GlyRs containing more than one β subunit should be impaired in Cl^-^ conduction. A hypothetical model of (α2)_3_β_2_ was constructed based on our open/desensitized structures using a α2-β-α2-β-α2 (meta) configuration where two conduction barriers formed (Figure 4E). First, negative electrostatic potentials in the pore near β:E297 and β:E290 indicate greatly reduced local Cl^-^ concentration as well as raised electrostatic barrier. Second, the two β:F282 sidechains resulted in tight constriction (<1.8Å) even in the open/desensitized states. To test whether heteromeric GlyR with more than one β subunit is indeed deficient in Cl^-^ conduction, we generated a chimeric construct, βchim, by combining α2-ECD and β-TMD, to overcome the invariant 4:1 stoichiometry in wild-type α2β GlyR. The elution peak of α2:βchim exhibited a left-shifting trend as α2:βchim expression ratios changed from 1:3 (Figure 4F solid blue) to 1:10 (Figure 4F solid red), and to 0:1 (Figure 4F solid green), suggesting arbitrary α2:βchim stoichiometry and homomeric βchim GlyR formation.

Although α2 and βchim formed GlyR with arbitrary stoichiometry biochemically, only the 4:1 configuration conducted Cl^-^. First, the amplitudes of glycine-activated currents decreased as more βchim is expressed with respect to α2 (Figure 4G, 1:3 and 1:10). When only the βchim is expressed, glycine elicited no current (Figure 4G, 0:1). The reduction in current is a result of GlyR being non-conductive when 2 or more βchim are incorporated, for the following reasons. First, small change in expression levels (<4 folds, Figure 4F) does not account for decimated current levels (>1000 folds). Second, PTX affinities stayed the same as native heteromeric GlyRs in all expression levels (Figure 4H). Since the incorporation of each βchim contributes equal amount of steric hindrance at β:F282 to destabilizing PTX binding, a same PTX affinity indicates a fixed number (1) of β subunit in each conductive GlyR. Higher βchim expression levels drive the population of the only conductive species, 4:1 α2:βchim GlyR, to other stoichiometries, therefore decreasing total measurable current. The “dominant negative” effect reported earlier of a similar chimeric β construct also supports this conclusion (Kuhse et al., 1993).

### α2β GlyR distribution in the cellular membrane

Single-molecule counting allows quantification of membrane proteins in native cell membranes (Patrizio and Specht, 2016; Schmidt et al., 1996). Most research using this method suggested 2 or 3 β subunits in each α1β and α3β GlyRs (Durisic et al., 2012; Patrizio et al., 2017). Since we have already demonstrated an invariant 4:1 α2:β stoichiometry for the α2β GlyR, this method allowed us to understand how α2β GlyR is distributed in cell membranes.

One or two homomeric α2 GlyRs were observed in each diffraction limited fluorescent spot (∼250 nm). We used an α2-GFP fusion (see methods), which is known to form pentameric GlyR (Du et al., 2015; Huang et al., 2015; Langosch et al., 1988), to determine the probability, *p_m_*, of GFP being mature and fluorescent under our expression conditions. In 2295 diffraction-limited fluorescent spots, many had photobleaching steps ranging from 1 to 5 (Figure 5, A and C) while a significant number had between 6 and 10 steps (Figure 5, B and C), suggesting the presence of two α2 GlyRs in some spots. Using a dual-binomial model in data fitting (Figure 5C), we derived a *p_m_* of ∼62%, with 67% of the spots containing 1 and 33% of the spots containing 2 α2-GFP GlyRs. Coincident with our findings, multiple α1 GlyRs being present in a single spot has also been reported (McGuire et al., 2012).

**Figure 5.**
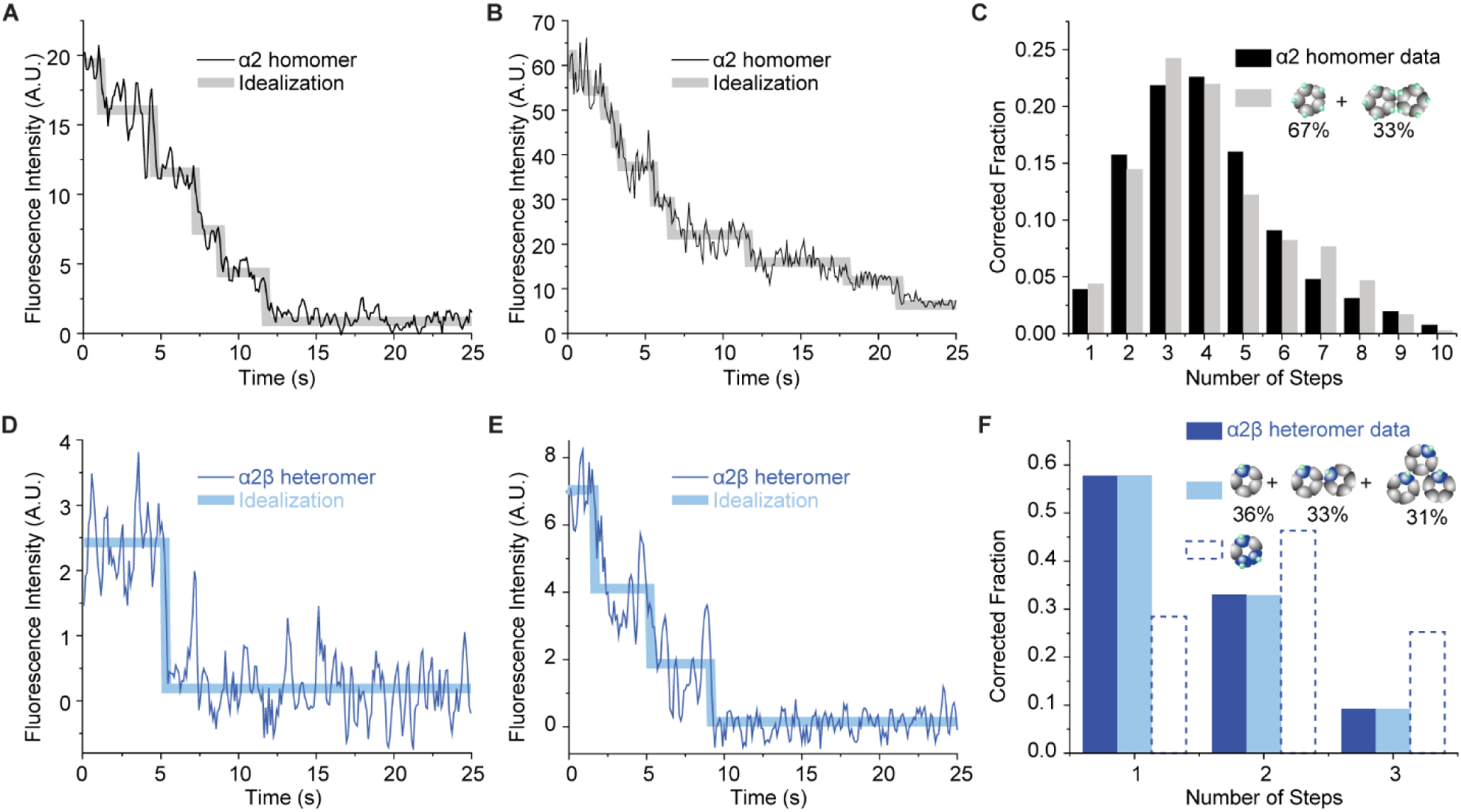
**Step-wise photobleaching of GlyRs in HEK293 cells.** (A and B) Representative fluorescence intensities (black) and idealization (grey) of α2-GFP exhibiting (A) 5 and (B) 10 photobleaching steps. (C) Histogram of the α2-GFP GlyR steps from 2295 spots after correction for missed events (black). A dual-binomial model was used to fit the data (grey), resulting in a GFP maturation rate pm of 62%. (D and E) Representative photobleaching traces (blue) and idealization (cyan) of α2β with (D) 1 and (E) 3 steps. (F) Histogram of the α2β GlyR steps (blue) from 206 spots, shown with two binomial models (cyan and dashed) using pm of 62%.

Up to three α2β GlyRs were found residing within diffraction limited distance. When α2β GlyR was expressed with GFP fusion in only the β subunit, bleaching steps ranged from 1 to 3 (Figure 5, D-F). Since each α2β GlyR contains only 1 β subunit, 3 bleaching steps necessitates no less than 3 α2β GlyR within diffraction limited distance. Indeed, using a tri-binomial model with *p_m_* fixed at 62%, we derived the fraction of spots containing 1, 2 and 3 α2β GlyRs to be 36%, 33% and 31%, respectively (Figure 5F light blue, see methods). As expected, our data cannot be explained with only one (α2)_2_β_3_ GlyR being in each spot (Figure 5F, dashed lines).

## DISCUSSION

Heteromeric GlyRs are the predominant form of GlyRs in adult animals. Research on embryonic/ early developmental forms of GlyRs, the homomeric GlyRs, provided very little information on how adult heteromeric GlyRs work. Even the subunit stoichiometry was disputed for decades (Burzomato et al., 2003; Durisic et al., 2012; Grudzinska et al., 2005; Kuhse et al., 1993; Patrizio et al., 2017; Yang et al., 2012). In this work, we developed a system that allowed characterization of heteromeric GlyRs both in cells and *in vitro*. Our results showed unequivocally that each physiological α2β GlyR contains only one β subunit and revealed the molecular mechanism underlying such arrangement. In addition, we identified structural features that gave rise to the specific electrophysiological properties, including glycine activation, single-channel conductance and picrotoxin blocking, of heteromeric GlyRs and showed why GlyRs containing more than 1 β subunit do not conduct Cl^-^. We further found in cellular membranes that multiple GlyRs may exist within diffraction limited distance (∼250 nm). Through these findings, we pin-pointed the only physiological form of α2β GlyR that conducts Cl^-^, and revealed its working mechanism.

Clustering of GlyR at post-synaptic densities is crucial for it function in neurotransmission. Since GlyR clustering relies on a specific interaction between the β subunit and the scaffolding protein gephyrin (Kirsch et al., 1993; Meyer et al., 1995), little can be learned from homomeric GlyRs since they do not contain the β subunit. Our results have strong implications in GlyR clustering as follows (Figure 6). Since gephyrin assembles into a multimeric form with one GlyR β subunit binding site in each monomer, the number of β subunits in each GlyR determines how GlyRs are clustered – having only 1 β subunit requires the gephyrin to cluster by itself, GlyRs are recruited to the already formed gephyrin clusters. Having more than 1 β subunit in each GlyR would allow multi-valent interactions between the gephyrin and GlyR, inducing cluster formation. This is reminiscent of the phase-separated systems where some factors serve as scaffolds and others as clients (Banani et al., 2016) – the 4:1 α2:β GlyR can only function as clients (Figure 6A) whereas additional β subunits would allow GlyR to act as scaffolds of the inhibitory post-synaptic density (Figure 6B). Our data shows that functional GlyRs can only serve as clients, indicating that synaptic plasticity of a given glycinergic synapse may be regulated through two mechanisms: the physical properties of post-synaptic density, for examples size and compactness, can be regulated through gephyrin, while the intensity of glycinergic transmission can be further tuned through GlyRs diffusion in and out of clusters. Clearly, more research is required to gain complete understanding.

**Figure 6.**
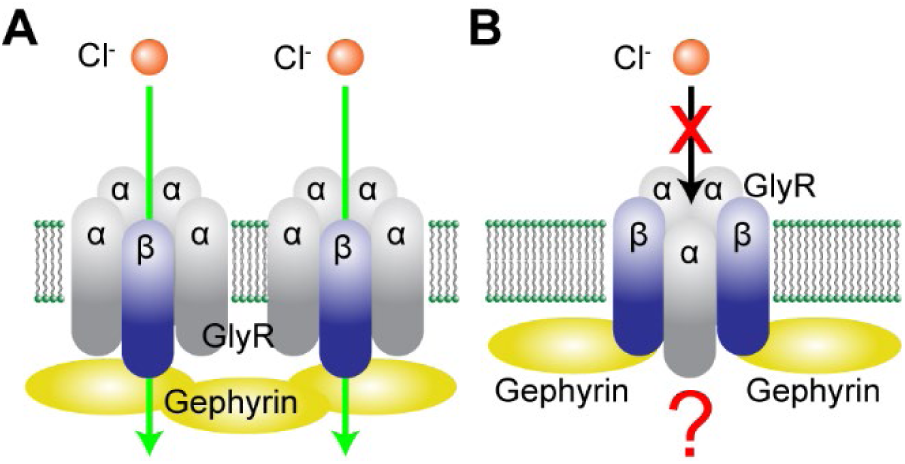
**Models of heteromeric GlyR – gephyrin clustering.** (A) GlyRs containing only one β subunit conducts Cl^-^ upon activation and interact with already-formed gephyrin clusters in a monovalent manner. (B) GlyRs containing two or more β subunits may form multivalent interactions with gephyrin but do not conduct Cl^-^.

Our results do not exclude the possibility of alternative subunit composition in native neuronal cells. First, we focused on the α2β GlyR, in contrast to α1β and/or α3β GlyR reported previously, raising the possibility that 4:1 is specific to the α2β GlyR. However, this seems unlikely because of the high sequence homology at the N-terminus and subunit interfaces (Figure S5 and S6). Second, unidentified chaperons and other factors might produce GlyRs with multiple β subunit in specific neuronal cells. In this case, since GlyRs containing multiple β subunits are non-conductive upon glycine-activation (Figure 4), these GlyRs will not function as ionotropic receptors, but instead as scaffolding proteins (Figure 6B). Multiple GlyRs residing within one diffraction-limited spot (∼250 nm in diameter) indicates that the number of β subunits in each spot do not necessarily equal the number of β subunits in each GlyR. Since we found this for both homomeric and heteromeric GlyRs, and in the absence of over-expressed gephyrin, the mechanism and physiological significance of this phenomenon calls for further investigations.

## ACKNOWLEDGMENTS

We thank Robbie Boyed for preparing tissue culture, molecular biology and protein biochemistry tools for this project; Dr. Khuloud Jaqaman, Dr. Xian Cheng and Dr. Bruno Rocha Azevedo for help in photobleaching data collection and analysis; Dr. Ryan Hibbs and Jeong Joo Kim for input to biochemistry experiments and comments on the manuscript; Dr. Roderick MacKinnon, Dr. Youxing Jiang and Dr. Xiao Tao for comments on the manuscript; All members of the Wang laboratory for helpful discussion. Cryo-EM data were collected at the University of Texas Southwestern Medical Center Cryo-EM Facility, which is funded by the CPRIT Core Facility Support Award RP170644.

## AUTHOR CONTRIBUTIONS

H.Y. performed the experiments and analyzed the data. X.B. collected the Cryo-EM data and reconstructed density maps. W.W. conceived the project, performed experiments, and analyzed data. W.W., H.Y. and X.B. composed the manuscript.

## METHODS

### Cloning of hGlyR constructs

DNAs coding human glycine receptor α2 (NCBI: NP_002054.1) and β (NCBI: NP_000815.1) were amplified from cDNA clones (McDermott Center, UT Southwestern Medical center). The amino acid numbering starts from the first amino acid after signal peptide. For GlyR α2 subunit, region encoding M3/M4 loop (residues Q317-K381) was substituted by GSSG peptide, this construct was named as α2em. The βem construct contained the following modifications: region encoding M3/M4 loop (residues N334-N377) was removed and GGSSAAA-monomeric enhanced green fluorescent protein (mEGFP)(Zacharias et al., 2002) -SGSGSG was inserted. A PA-tag (GVAMPGAEDDVV)(Fujii et al., 2014) and PreScission protease site (LEVLFQ/GP)(Walker et al., 1994) were inserted after signal peptide at the N-terminus. For GlyR α2-Tsi-GFP construct, region encoding M3/M4 loop (residues Q317-K381) was replaced with GGSSAAA-Tsi3(Li et al., 2013)-GSAGSAAGSG-mEGFP-SGSGSG. These mutants were cloned into a modified BacMam expression vector(Goehring et al., 2014). α2 wild type and α2em were cloned into pLVX-IRES-ZsGreen1 vector (Clonetech) for electrophysiology. The α2em and βem constructs are used throughout this work and denoted α2 and β for simplicity unless otherwise specified.

### Fluorescence-Detection Size-Exclusion Chromatography (FSEC) expression assay

The BacMam vectors containing target protein were transformed into DH10BacY competent cells (Geneva Biotech) to produce bacmids, which in turn were used to produce viruses in *sf9* insect cells. The virus titer was determined using published methods(Goehring et al., 2014; Morales-Perez et al., 2016) and the virus ratios in this work denotes the multiplicity of infection (MOI) ratios. HEK293T adherent cells were infected with viruses at different α2:β ratio or α2:α2-Tsi-GFP ratios (1:0, 20:1, 10:1, 5:1, 3:1, 1:1, 1:3, 1:5, 1:10, 1:20, 0:1, total virus amount was kept constant). Expression was induced 20 hours after infection by adding 10 mM sodium butyrate with 5μM strychnine added at the same time. Cells were collected 48 hours after induction by centrifugation at 20,000 g for 30 minutes at 4°C. Cell pellets were resuspended and extracted with buffer A (40 mM Tris pH 8.0, 200 mM NaCl, 0.2 mM PMSF, 0.75%(w/v) n-Dodecyl β-D-maltoside (DDM) and 0.075% (w/v) Na Cholate) for 1 hour at 4°C. After centrifuge, the supernatants were analyzed by FSEC on high-performance liquid chromatography system (HPLC, Shimadzu) equipped with fluorescence detectors set to read GFP fluorescence (excitation wavelength 480mm, emission wavelength 515mm). Due to differences in molecular weight (α2 45KDa, β 77KDa, α2-Tsi-GFP 87KDa), elution volume was a function of subunit stoichiometry.

### Expression and purification of hGlyR

Large scale expression of hGlyR was essentially the same as in FSEC assays except for using HEK293S GnTI^-^ cells in suspension culture instead of adherent in FSEC. HEK293S GnTI^-^ cells were infected at a density of 2.5 × 10^6^ cells/ml using viruses encoding GlyR α2 and GlyR β at 1:3 ratio. Expression was induced 20 hours after infection with 10 mM sodium butyrate and 5μM strychnine (or no strychnine for GlyR-glycine complex) added at the same time. 48 hours later, cells were collected by centrifugation, then homogenized and extracted with buffer A for 1 hour at 4°C. PA-tag antibody (NZ-1)(Fujii et al., 2014) resin was used to capture hGlyR heteromer from the extracted supernatant and then washed with buffer B (20 mM Tris pH 8.0, 200 mM NaCl, 0.2 mM PMSF, 0.1% DDM, 0.02% Na Cholate). PreScission protease was added to release the hGlyR heteromer from PA-tag antibody resin. Eluent was concentrated and loaded onto Superose 6 increase 10/300 GL size exclusion column (GE Healthcare) in buffer C (20 mM Tris pH 8.0, 200 mM NaCl, 0.025% DDM). Peak fractions were collected and concentrated to 4 mg/ml. GlyR α2-Tsi-GFP homomer was prepared the same way except for that the Tse3(Li et al., 2013) resin was used for purification. Buffer B containing 10 mM EDTA was used to elute hGlyR homomer from Tse3 resin. Concentrated proteins were used immediately.

### Structure determination using single-particle cryo-EM

#### Reconstitution of α2β heteromer into saposin nanodisc

Reconstitution of GlyR α2β heteromer into saposin nanodisc was modified from the published protocol(Frauenfeld et al., 2016). After screening, a 1:30:200 molar ratio of α2β:saposin:brain polar lipids extract (BPE) (Avanti) resulted in the best reconstitution with a small void peak corresponding to lipid vesicles without scaffold. 100μl α2β heteromer (4.5mg/ml) and 17μl BPE (13.6mg/ml, 1% DDM) were mixed and put at room temperature (RT) for 10 minutes. Then 393μl saposin(1.2mg/ml) was added and well mixed. The mixture was put at RT for 10 minutes, then 3.2mL dilution buffer (20mM Tris pH 8.0, 200mM NaCl, 1mM strychnine / 2mM glycine) was added. After several minutes, bio-beads SM-2 (Bio-Rad) were added to the mixture and rotated overnight at 4 °C. Next morning, the mixture was put into a new tube to remove the bio-beads, then fresh bio-beads were added again. After 8 hours, the mixture was centrifuged and loaded onto Superose 6 increase size exclusion column in SEC buffer (20mM Tris pH 8.0, 200mM NaCl, 1mM strychnine / 2mM glycine). The peak fractions containing α2β-saposin nanodisc were concentrated to 4 mg/mL and used for EM grids preparation immediately.

#### Cryo-EM sample preparation, data collection and image processing

Fluorinated fos-choline 8 (Anatrace) was added into 4 mg/ml α2β heteromer saposin nanodisc sample at a final concentration of 1 cmc just before freezing grids. 3.5 µl sample applied to a glow-discharged Quantifoil R1.2/1.3 400-mesh gold holey carbon grid (Quantifoil, Micro Tools GmbH, Germany), blotted under 100% humidity at 4 °C and plunged into liquid ethane using a Mark IV Vitrobot (FEI). Micrographs were collected on a Titan Krios microscope (FEI) with a K3 Summit direct electron detector (Gatan) operating at 300 kV using the SerialEM software. The GIF-Quantum energy filter was set to a slit width of 20 eV. Images were recorded in the super-resolution counting mode with the pixel size of 0.83 Å. Micrographs were dose-fractioned into 32 (strychnine complex) or 50 (glycine complex) frames with a dose rate of 1.6 e^−^/Å/frame. Motioncorr2 program(Zheng et al., 2017) was used to perform 2-fold binning, motion correction and dose weighting of the movie frames. CTF correction were carried out using the GCTF program(Zhang, 2016). The following image processing steps were carried out in RELION 3.0(Scheres, 2012). Auto-picked particles were extracted and binned by 4 times and used in 2D classification. Particles in good 2D classes were selected and subjected to 3D classification using an initial model downloaded from EMDB database (EMD-6346)(Du et al., 2015). 3D classification into 6 classes yielded 2 classes with good density for the entire channel. The cryo-EM density of one GFP were identified in both classes, but present at different position with respect to the channel. As GFP is only fused to the β subunit of the receptor, this 3D classification result suggests that there is only one β subunit but four α subunits in the hetero-pentameric channel, and the relatively position of this single β subunit with respect to the other four α subunits are different between these two good 3D classes. To put the β subunit of both classes in the same position that would allow us to combine these two good classes, we rotated each particle from one class around the Z axis by 360°/5 through manually modifying the column “_rlnAngleRot” in the RELION selection star file. As a result, a total of 111,378 particles for GlyR-strychnine and 268,380 particles for GlyR-glycine complexes from two good 3D classes were combined, re-extracted to the original pixel size. 3D refinement of these particles with 8° local angular search and the C1 symmetry imposed led to a 3D reconstruction to overall resolutions of 3.8 – 3.9 Å, although the transmembrane region was resolved at relatively lower resolution. Therefore, the particles of the initial 3D reconstruction were subjected to partial signal subtraction to keep only the transmembrane region(Bai et al., 2015). 3D classification of these partial-subtracted particles without alignment led to two classes that showed much better density for the transmembrane domain (Figure S2A). After reverting to un-subtracted version, CTF refinement, particle polishing, and refinement using C1 symmetry, GlyR-strychnine complex yielded two classes. One class contained 38,541 particles and reached an overall resolution of 3.8 Å (closed 1) and the other contained 44,965 particles and reached 3.6 Å (closed 2). Closed 2 class exhibited a more 5-fold symmetrical TM arrangement (Figure S5). GlyR-glycine complex also yielded two classes, one contained 55,816 particles with 3.8 Å (open) resolution and the other contained 56,419 particles and reached 3.6 Å (desensitized) resolution. Resolutions were estimated by applying a soft mask around the protein densities with the Fourier Shell Correlation (FCS) 0.143 criterion. Local resolutions were calculated using Resmap(Kucukelbir et al., 2014).

#### Model building and refinement

Model building of GlyR α2β heteromer was initiated by docking the structure of α1 homomer strychnine-bound state(Du et al., 2015) (PDB ID: 3JAD) into the density map of GlyR α2β heteromer using Chimera(Pettersen et al., 2004). Amino acids were mutated according to α2 and β sequences and rebuilt in Coot(Emsley et al., 2010). The structure model was manually adjusted in Coot, then refinement was performed with real-space refinement module with secondary structure restraints in PHENIX package(Adams et al., 2010; Liebschner et al., 2019). MolProbity was used to validate the geometries of the refined models(Williams et al., 2018). Fourier shell correlation (FSC) curves were calculated between refined atomic model and the work/free half maps as well as the full map to assess the correlation between the model and density map. Statistics of cryo-EM data processing and model refinement are listed in Table S1. Pore radii were calculated using the HOLE program(Smart et al., 1996). Figures were prepared in Chimera or PyMOL(Schrödinger, 2015). Electrostatics were calculated using Adaptive Poisson-Boltzmann plugin at 37°C and with 150 mM of monovalent anion/cations (Baker et al., 2001). The final model of strychnine closed 2 contained the α2 and β subunit amino acids except the following: α2 subunit (total 364aa, 339aa built, 25aa not built) K1-T14, GSSG linker, K382 and E420-K425; β subunit (total 444aa, 338aa built, 106aa not built) K1-Q20, L332-N333, GGSSAAA-EGFP-SGSGSG insertion and V378-A448. Q21-R28 of β subunit were modeled as poly-alanine due to limit in resolution. The model for strychnine closed 1 is the same as closed 2 except for Q21-R28 of β subunit were not modeled. The final model of glycine bound desensitized state contained the α2 and β subunit amino acids except the following: α2 subunit (total 364aa, 338aa built, 26aa not built) K1-T14, GSSG linker, K382 and H419-K425; β subunit (total 444aa, 337aa built, 107aa not built) K1-R28, GGSSAAA-EGFP-SGSGSG insertion and V378-V443. The model for glycine bound open state is the same as desensitized state except that only 3 glycine are modeled.

### Electrophysiology

#### Whole cell patch clamp

Plasmids bearing GlyR constructs were transiently transfected into HEK293T cells by Lipofectamine 3000 (Invitrogen) following manufacturer’s protocol. For each 35 mm dish, 1.5 μg of DNA was transfected at different α:β ratios (1:0, 1:1, 1:3, 1:10). After transfection, cells were cultured at 37°C for 6∼8 hours, then transferred to 30°C cultured for ∼12 hours. Whole-cell recordings were performed at room-temperature. The bath solution contained (in mM): 10 HEPES pH 7.4, 10 KCl, 125 NaCl, 2 MgCl_2_, 1 CaCl_2_ and 10 glucose. The pipette solution contained (in mM): 10 HEPES pH 7.4, 150 KCl, 5 NaCl, 2 MgCl_2_, 1 CaCl_2_ and 5 EGTA. Borosilicate glass pipettes with resistance between 2∼7 MΩ were used. A Digidata 1550B digitizer (Molecular Devices) was connected to an Axopatch 200B amplifier (Molecular Devices) for data acquisition. Analog signals were filtered at 1 kHz and subsequently sampled at 20 kHz and stored on a computer running pClamp 10.5 software. Membrane was held at −80 mV during perfusion of ligands to record GlyR current. For the heteromer recordings, as the GlyR β construct had an EGFP insertion, we used GFP fluorescence to identify the cells expressing the GlyR β subunit. Hill equation was used to fit the dose-response data and derive the EC_50_ (*k*) and Hill coefficient (*n*). For glycine activation, we used 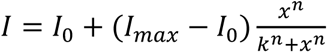, where *I* is current, *I_0_* is the basal current (spontaneous opening current and leak, very close to 0), *I_max_* is the maximum current and *x* is glycine concentration. Hill equation was also used to fit picrotoxin inhibition data: 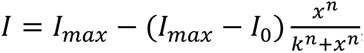, with same definitions of symbols except that *k* represents IC_50_. Data was fitted with software OriginPro (OriginLab, Northampton, MA). Measurements were from 3–6 cells, average and S.E.M. values were calculated for each data point.

#### Proteoliposome reconstitution

Purified α2β heteromer and α2 homomer were reconstituted into proteoliposomes using the lipid mixture composed of 3:1 (wt:wt) 1-palmitoyl-2-oleoyl-sn-glycero-3-phosphoethanolamine (POPE) : 1-palmitoyl-2-oleoyl-sn-glycero-3-phospho-1′-rac-glycerol) (POPG). The reconstitution procedure is similar to that previously reported(Wang et al., 2014). Briefly, 20 mg/ml of the above lipid mixture was dispersed by sonication until translucent and then solubilized with 1% DDM. GlyR was diluted with reconstitution buffer (10 mM potassium phosphate pH 7.4, 150 mM KCl, 1 mM EDTA and 3 mM DTT) supplemented with 0.2% DDM to 2 mg/ml. Equal volumes of 20 mg/ml solubilized lipid mixture was combined with the diluted GlyR solutions, resulting in a final protein:lipid (wt:wt) ratios of 1:10 at a lipid concentration of ∼10 mg/ml. Detergent was removed by dialysis against reconstitution buffer for 4 times, 12 hours each at 4°C. Bio-Beads SM2 (Bio-Rad) were added outside the dialysis bag (∼1 g/500mL buffer) in the last three cycles. 20 μL aliquots of the proteoliposomes were frozen stored at −80 °C until needed.

#### Planar lipid bilayer recording

The planer bilayer recording experiments are similar to previously described(Wang et al., 2016; Wang et al., 2014). Briefly, 20 mg/ml of lipid mixture composed of 2:1:1(wt:wt:wt) of DOPE:POPC:POPS, or brain polar extract (Avanti), was dissolved in decane and used to form planar lipid bilayer over a ∼100 μm hole on a piece of transparency film using the painting method. The same solution (10 mM potassium phosphate pH 7.4, 150 mM KCl, 1 mM EDTA) was used in both chambers. Proteoliposomes were mixed with 1M KCl at equal volume and sonicated to facilitate vesicle fusion into planer bilayer immediately before use. After vesicle fusion into planer bilayer, buffer containing glycine or strychnine was applied on the planar bilayer manually. A Digidata 1550B digitizer (Molecular Devices) interfaced to pClamp10.5 software was connected to an Axopatch 200B amplifier (Molecular Devices) for data acquisition. Analog signals were filtered at 1 kHz and subsequently sampled at 20 kHz. Recordings were performed at room temperature. Membrane voltage was held at −50 mV. The concentrations of glycine and strychnine applied to the membrane were 200 μM and 10 μM, respectively.

### Step-wise Photobleaching

α2-GFP or α2:β at 1:3 ratio was transfected into HEK293T cells on collagen coated glass bottom dishes using lipofectamine 3000 (Invitrogen) for expression of α2 GlyR or α2β GlyR, respectively. The transfected cells were kept in CO_2_ incubator for 6 hours at 37°C and then removed to 30°C for overnight expression. Cells were unroofed(Usukura et al., 2016) and fixed prior to imaging as follows: After 3 times wash at room temperature (RT) with Phosphate-Buffered Saline PBS (10 mM Na_2_HPO_4_, 1.8 mM NaH_2_PO_4_, 137 mM NaCl, 3 mM KCl), cells were sonicated at 20% power for 1 s using a Branson SFX550 sonicator equipped with a flat microtip in ice-cold PBS supplemented with 2% para-formaldehyde (PFA), 2 mM MgCl_2_ and 1 mM CaCl_2_. Unroofed cells were washed quickly with ice-cold PBS and immediately switched into PBS containing 4% PFA at RT and fixed for 15min. After 3 times washing in PBS for 10 min each, the cells were imaged within 2 hours. Total internal reflection illumination was achieved using an in-house built prism-type system using the Gem 488 laser (Laser Quantum) and components from Thorlabs at 10% power. A 1.5 OD neutral density filter was used during searching of cells to minimize photobleaching. A Leica DM6 FS microscope equipped with a 63x 1.2 NA CS2 water immersion objective and a Hamamatsu flash 4.0 V3 camera was used for imaging. Movies were collected using a PC running Metamorph (Molecular Devices) with 100 ms exposure for each frame and total exposure of 50 s. Regions of interest were subsequently cropped using ImageJ(Schneider et al., 2012) and analyzed using the PIF program(McGuire et al., 2012) to generate statistics of photobleaching steps. A dual-binomial model formulated allowing one and two pentameric GlyRs in each diffraction-limited fluorescence spot was used to fit the photobleaching step distribution of homomeric α2G GlyR as follows:

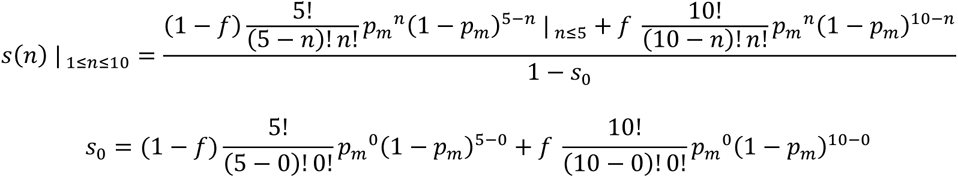

Where *s*(*n*) is the fraction of spots with *n* photobleaching steps corrected for the fraction of non-fluorescent events *s_0_*. *p_m_* and *f* were the fitted parameters, which denote the maturation rate of GFP and the fraction of spots with two α2 GlyRs, respectively. Similarly, a tri-binomial model assuming simultaneous presence of 1, 2 and 3 GlyRs in each spot was used for the heteromeric α2β GlyR photobleaching steps as follows

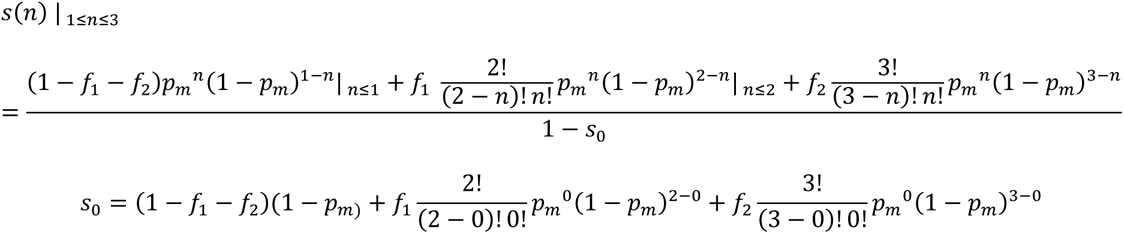

Where *f_1_* and *f_2_* were the fitted parameters which denote the corrected fractions of spots with 2 and 3 α2β GlyRs, respectively. *p_m_* was fixed at 0.62 (the value from α2 GlyR). Traces and statistics were plotted using OriginPro (OriginLab).

### Western Blotting

For GlyR α2 subunit, a V5-tag (KEGKPIPNPLLGLDSTAA) was inserted into the α2em construct after signal peptide at the N-terminus. Plasmids bearing GlyR constructs were transiently transfected into HEK293T cells by Lipofectamine 3000 (Invitrogen) following manufacturer’s protocol at different α2:β DNA ratios (1:0, 3:1, 1:1, 1:3, 1:10, 0:1). After transfection, cells were cultured at 37°C for 20 hours, then transferred to 30°C for additional ∼48 hours. After detergent extraction, supernatants were mixed with an equal volume of loading buffer containing 4% SDS and 10% DTT. Then samples were analyzed using SDS-PAGE with Bio-Rad 4-15% TGX gels and transferred onto nitrocellulose membrane. Western Blot was performed with anti-V5-tag (Cell Signaling Technology, D3H8Q) for α2 or Anti-GFP (Santa Cruz Biotechnology, sc-9996) for β.

## DATA AND SOFTWARE AVAILABILITY

Data Resources

Atomic coordinates of α2β glycine receptor have been deposited in the Protein Data Bank with accession numbers 7KUY for GlyR α2β closed state 2, 7L31 for GlyR closed state 1, 5BKG for glycine bound open state and 5BKF for glycine bound desensitized state. The corresponding cryo-EM density maps have been deposited with the Electron Microscopy Data Bank using accession numbers EMD-23041 for α2β closed state 2 and EMD-23148 for α2β closed state 1, EMD-9404 for glycine bound open state and EMD-9403 for glycine bound desensitized state.

---

**Figure S1.**
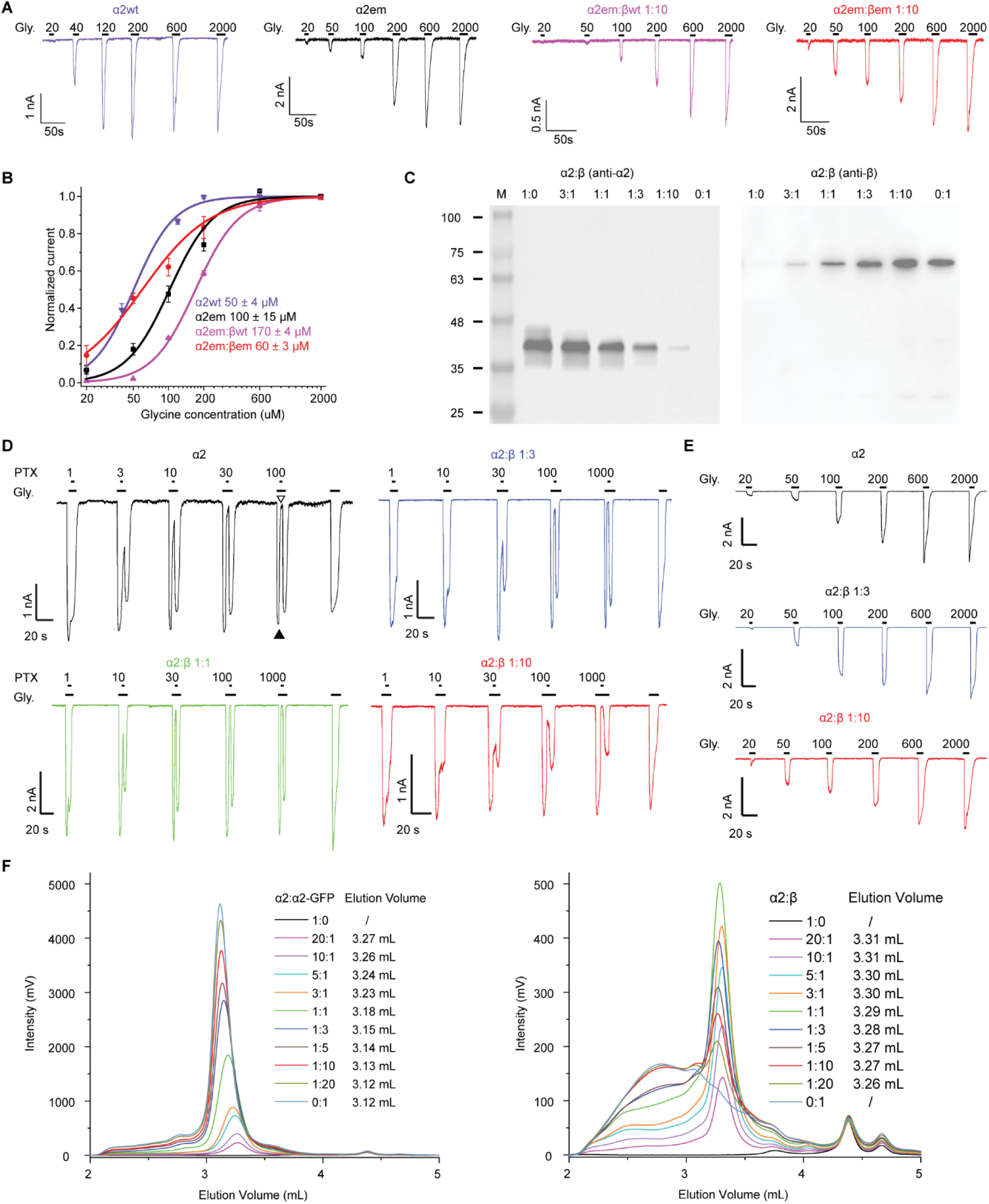
**Whole cell voltage-clamp electrophysiology and expression assays of α2 and β GlyRs constructs, related to Figure 1.** (A) Representative voltage-clamp recordings of glycine dose response in HEK293 cells transfected with α2wt (magenta), α2em (black), α2em:βwt 1:10 (pink) and α2em:βem 1:10 (red) DNAs. (B) Dose response of glycine from (A). Data points with S.E.M. (n = 3-6 cells) were plotted with Hill fits and colored the same as in (A). EC50 values are listed. (C) Western blot of α2 and β subunits at different DNA ratios. M: molecular weight marker, molecular weights are in kDa. See methods for details. (D) PTX inhibition in the presence of 100 μM glycine. α2:β expression ratios of 1:0, 1:1, 1:3 and 1:10 were used. The PTX and glycine concentrations are in μM. (E) Glycine activation at α2:β ratios of 1:0, 1:3 and 1:10. The glycine concentrations are in μM. (F) FSEC of α2 and α2β GlyRs at different α2:β virus ratios. Left panel, FSEC of α2 GlyR expressed at listed α2:α2-Tsi-GFP virus ratios. Right panel, FSEC of α2β GlyR expressed at different α2:β virus ratios. Elution volumes are listed beside each virus ratio. Molecular weight of α2, α2-Tsi-GFP and β are 45KDa, 87KDa and 77KDa, respectively.

**Figure S2.**
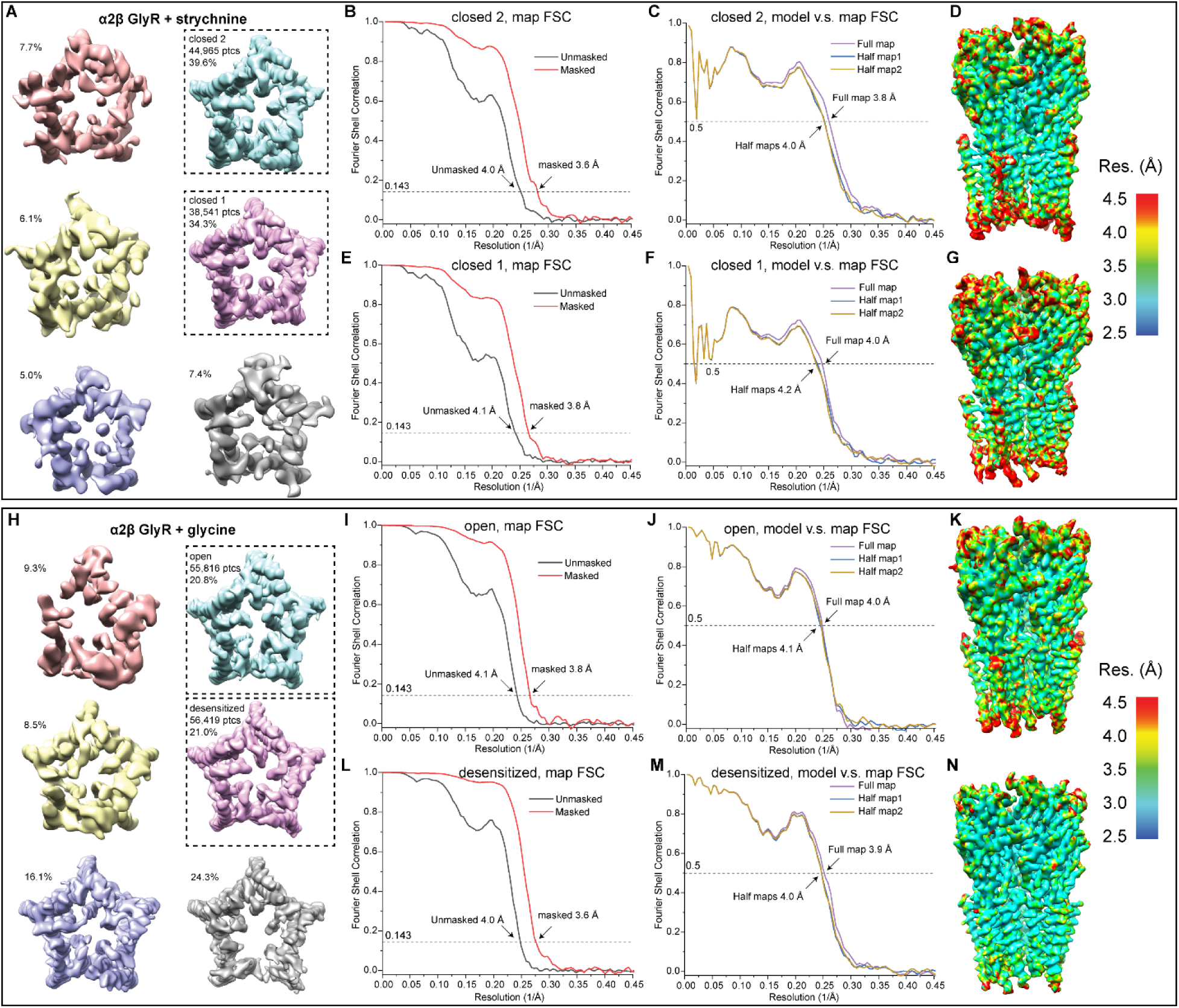
**Classes of α2β GlyR density maps, related to Figure 3.** (A-G) α2β GlyR closed states. (A) 6 classes resulting from 3D classification after signal-subtraction view along the ion conduction pore from the intracellular side. The percentages of particles in each class are indicated. The two good classes were subsequently refined. (B) The gold standard Fourier Shell Correlation (FSC) curves from the final 3D refinement of closed state 2, with (red) and without (black) mask. (C) FSC between the atomic model and half map 1 (blue), half map 2 (orange) and the full map (purple) of closed state 2. (D) Density map of closed state 2 colored according to its local resolution estimated using RESMAP. (E) The gold standard FSC from the final map of closed state 1. (F) FSC between the atomic model and maps of closed state 2. Color codes are the same as in (B) and (C). (G) Density map of closed state 1 colored according to its local resolution estimated using RESMAP. (H-N) α2β GlyR open state and desensitized state. (H) 6 classes resulting from 3D classification. The two good classes were subsequently refined. (I-J) FSC curves of open states. (K) Density map of open state colored according to its local resolution (L-M) FSC curves of desensitized states. (N) Density map of open state colored according to its local resolution.

**Figure S3.**
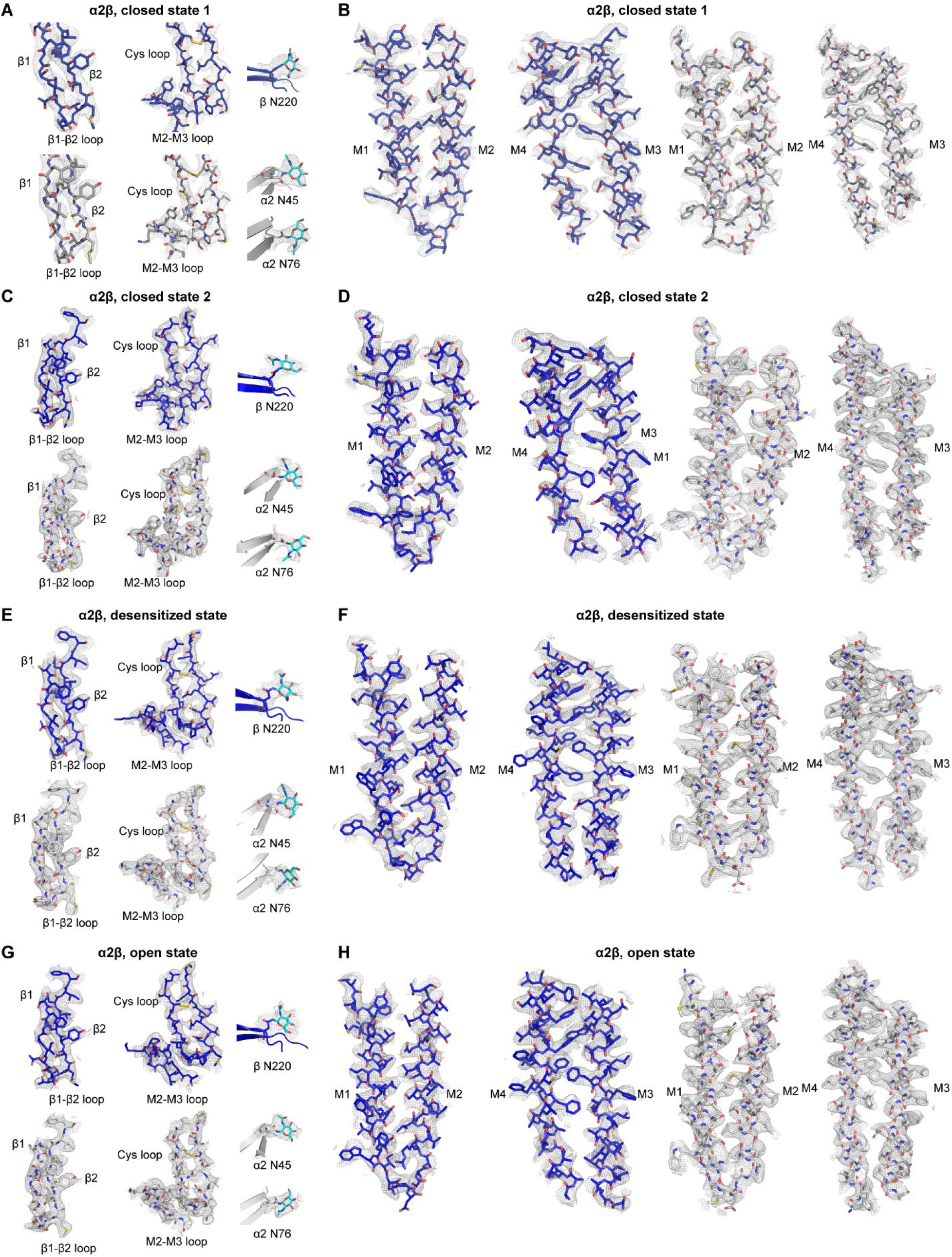
**Typical density maps, related to Figure 3 and Figure 4.** (A-B) Densities maps of representative regions of α2β GlyR closed state 1. (A) Density maps of β-sheet, Cys-loop and M2-M3 loop in β subunit ECD; N-glycosylation on the β subunit ECD; β-sheet, Cys-loop and M2-M3 loop in α2 subunit ECD; Two N-glycosylation sites on α2 subunit ECD. (B) M1-M4 Helices of β subunit and α2 subunit TMD. (C-D) Densities maps of representative regions of α2β GlyR closed state 2. Regions are the same as (A-B). (E-F) Densities maps of representative regions of α2β GlyR desensitized state. Regions are the same as (A-B). (G-H) Densities maps of representative regions of α2β GlyR open state. Regions are the same as (A-B).

**Figure S4.**
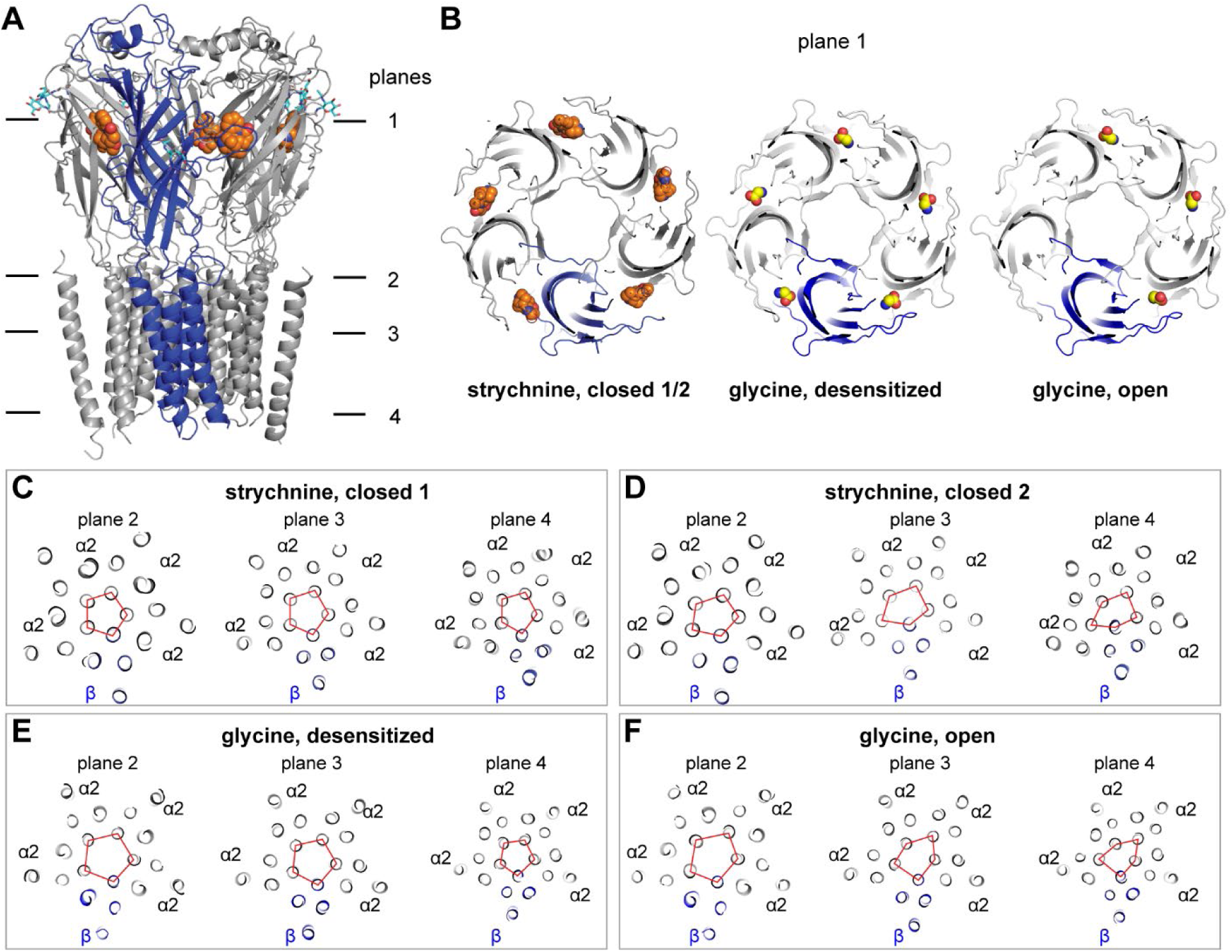
**Symmetry of the α2β GlyR, related to Figure 3.** (A) α2β GlyR closed state 2 viewed parallel to the membrane with four planes indicated. (B) Top views from the extracellular side clipped at plane 1 of closed, desensitized, and open states. (C) Top view of planes 2-4 for closed state 1. (D) Top view of planes 2-4 for closed state 2. (E) Top view of planes 2-4 for desensitized state. (F) Top view of planes 2-4 for open state. α2 subunit is colored in gray, β in blue, strychnine in orange as spheres, glycine in yellow as spheres and NAG in cyan as sticks. The apexes of the red pentagons in panels (C-F) indicate the M2 helices positions.

**Figure S5.**
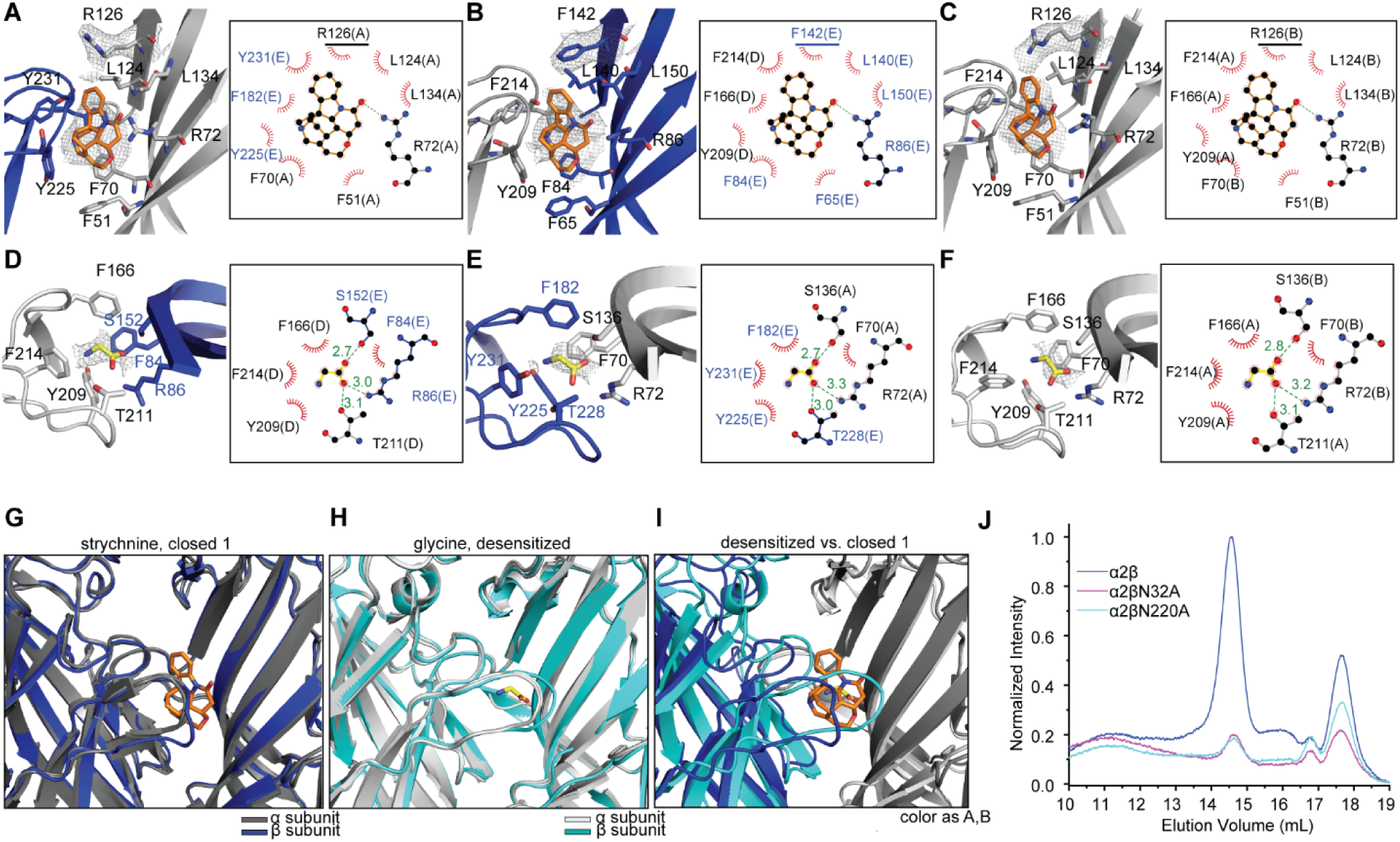
**Strychnine and glycine binding sites, related to Figure 3 and Figure 4.** (A-C) Strychnine binding sites between (A) β/α2, (B) α2/β and (C) α2/α2 interfaces from closed state 2. Densities of α2 R126, β F142 and Strychnine are contoured at 3σ. Strychnine (orange) and the residues interacting with strychnine (α2: grey, β: blue) are shown as sticks. LigPlot schematic of strychnine bind sites between (A) β/α2, (B) α2/β and (C) α2/α2 interfaces, showing electrostatic (dashes) and hydrophobic (eyelashes) interactions. Non-conservative residues α2 R126 and β F142 are underlined. (D-F) Glycine binding sites between (D) α2/β, (E) β/α2 and (F) α2/α2 interfaces from desensitized state. Densities of glycine are contoured at 4σ. Glycine (yellow) and the residues interacting with glycine (α2: grey, β: blue) are shown as sticks. LigPlot schematic of glycine bind sites between (D) α2/β, (E) β/α2 and (F) α2/α2 interfaces, showing electrostatic (dashes) and hydrophobic (eyelashes) interactions. (G) Strychnine binding sites between β/α2, α2/β and α2/α2 interfaces from closed state 1. Strychnine (carbon as orange, nitrogen as blue and oxygen as red) are shown as sticks, α2 and β (α2: grey, β: blue) are shown as cartoon. (H) Glycine binding sites between β/α2, α2/β and α2/α2 interfaces from desensitized state. Glycine (carbon as yellow, nitrogen as blue and oxygen as red) are shown as sticks, α2 and β (α2: grey, β: cyan) are shown as cartoon. (I) Overlap of glycine/strychnine binding site between β/α2 interface from desensitized/closed 1 state. The colors are same as (G) and (H). (J) FSEC of α2β GlyR with glycosylation site mutation. Wild type α2β are colored as blue, α2β with β:N32A mutation are colored as pink, α2β with β:N220A mutation are colored as cyan.

**Figure S6.**
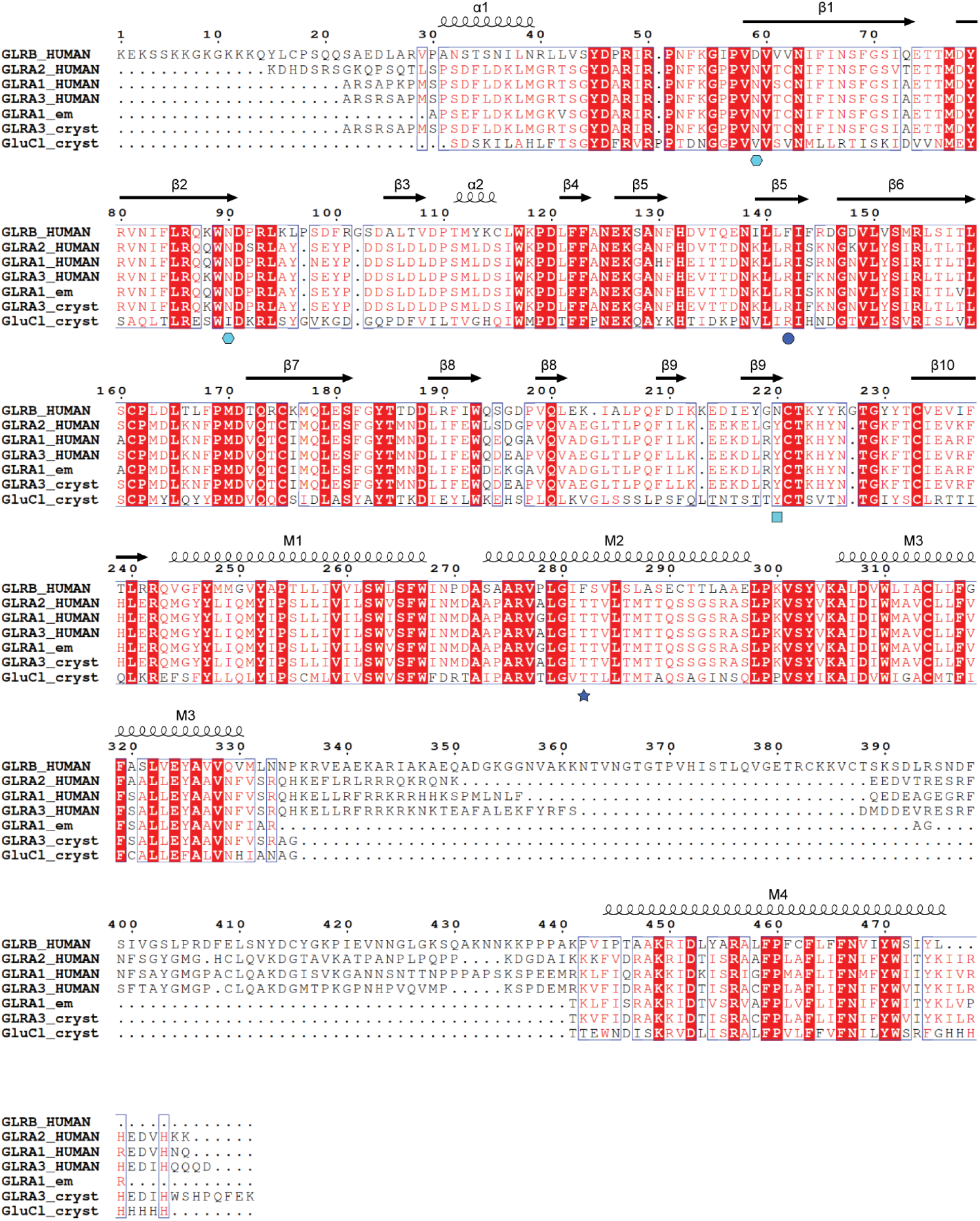
**Sequence alignment of GlyR subunits, related to Figure 4.** Signal peptides have been removed before sequence alignment. Residue conservation is indicated by blue rectangle and red highlights. Secondary structure elements are indicated by helices for α-helices and arrows for β-strands above the alignment. N-linked glycosylation site of GlyR α2 and β are indicated by cyan hexagon and cyan rectangle, respectively. Non-conservative residue between GlyR α2 and β involved in strychnine binding is indicated by blue circle. The Phe282 in M2 helix of GlyRβ is indicated by blue star. The alignment was generated by Clustal W. Protein sequences alignment: human GLRB (Uniprot P48167), human GLRA2 (Uniprot P23416), human GLRA1 (Uniprot P23415), human GLRA3 (Uniprot O75311), GLRA1em(PDB accession number 3JAE), GLRA3cryst (PDB accession number 5CFB) and GluClcryst (PDB accession number 4TNV).

**Table S1.**
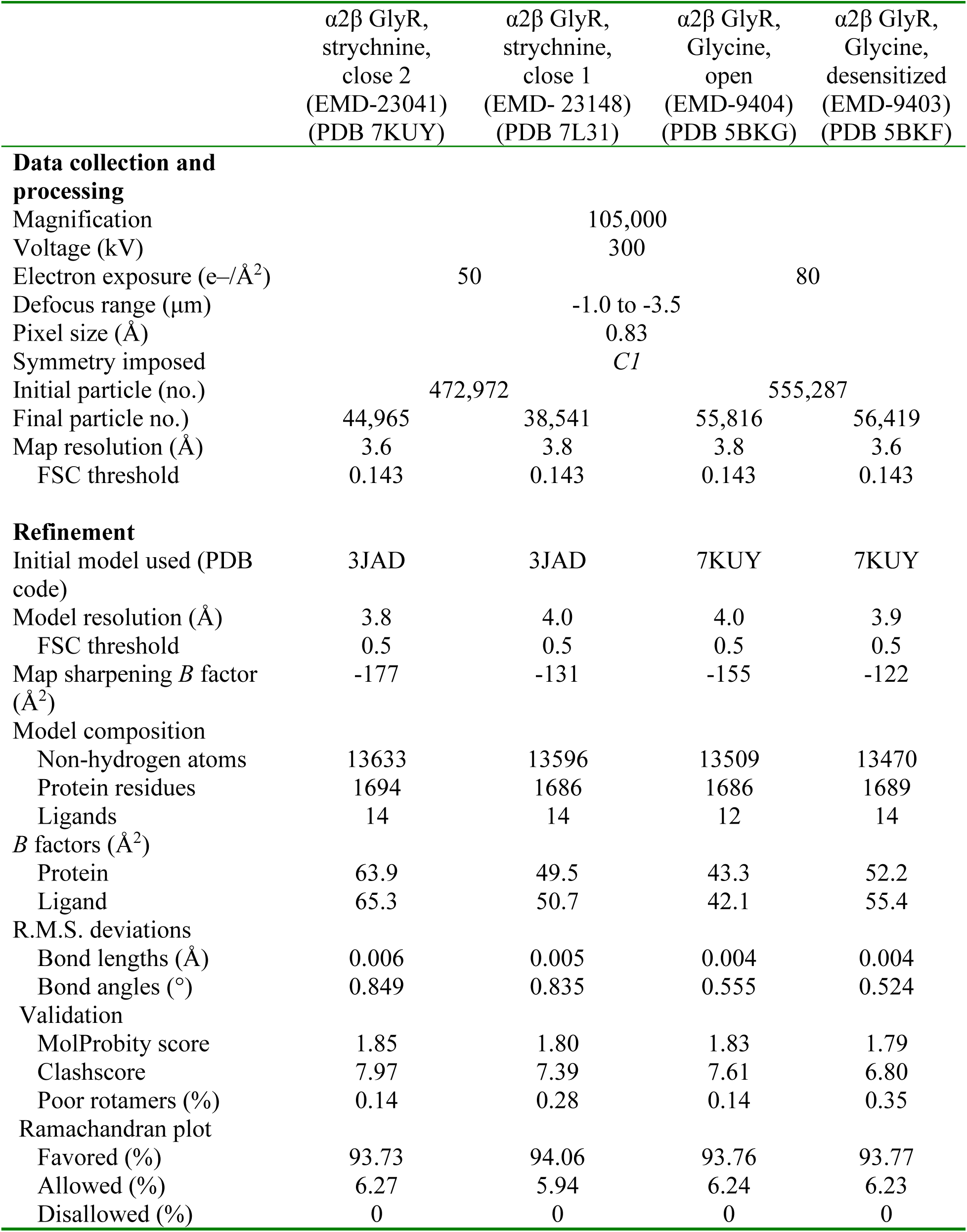
Cryo-EM data collection, refinement and validation statistics

## REFERENCES

Adams, P.D., Afonine, P.V., Bunkoczi, G., Chen, V.B., Davis, I.W., Echols, N., Headd, J.J., Hung, L.W., Kapral, G.J., Grosse-Kunstleve, R.W., et al. (2010). PHENIX: a comprehensive Python-based system for macromolecular structure solution. Acta Crystallogr D Biol Crystallogr 66, 213–221.

Ali, D.W., Drapeau, P., and Legendre, P. (2000). Development of spontaneous glycinergic currents in the Mauthner neuron of the zebrafish embryo. Journal of neurophysiology 84, 1726–1736.

Bai, X.C., Rajendra, E., Yang, G., Shi, Y., and Scheres, S.H. (2015). Sampling the conformational space of the catalytic subunit of human gamma-secretase. Elife 4.

Baker, N.A., Sept, D., Joseph, S., Holst, M.J., and McCammon, J.A. (2001). Electrostatics of nanosystems: application to microtubules and the ribosome. Proceedings of the National Academy of Sciences of the United States of America 98, 10037–10041.

Banani, S.F., Rice, A.M., Peeples, W.B., Lin, Y., Jain, S., Parker, R., and Rosen, M.K. (2016). Compositional Control of Phase-Separated Cellular Bodies. Cell 166, 651–663.

Becker, C.M., Hoch, W., and Betz, H. (1988). Glycine receptor heterogeneity in rat spinal cord during postnatal development. The EMBO journal 7, 3717–3726.

Bode, A., and Lynch, J.W. (2014). The impact of human hyperekplexia mutations on glycine receptor structure and function. Molecular brain 7, 2.

Bormann, J., Rundstrom, N., Betz, H., and Langosch, D. (1993). Residues within transmembrane segment M2 determine chloride conductance of glycine receptor homo- and hetero-oligomers. The EMBO journal 12, 3729–3737.

Burzomato, V., Groot-Kormelink, P.J., Sivilotti, L.G., and Beato, M. (2003). Stoichiometry of recombinant heteromeric glycine receptors revealed by a pore-lining region point mutation. Receptors Channels 9, 353–361.

Cascio, M., Schoppa, N.E., Grodzicki, R.L., Sigworth, F.J., and Fox, R.O. (1993). Functional expression and purification of a homomeric human alpha 1 glycine receptor in baculovirus-infected insect cells. The Journal of biological chemistry 268, 22135–22142.

Cascio, M., Shenkel, S., Grodzicki, R.L., Sigworth, F.J., and Fox, R.O. (2001). Functional reconstitution and characterization of recombinant human alpha 1-glycine receptors. The Journal of biological chemistry 276, 20981–20988.

Citri, A., and Malenka, R.C. (2008). Synaptic plasticity: multiple forms, functions, and mechanisms. Neuropsychopharmacology : official publication of the American College of Neuropsychopharmacology 33, 18–41.

Du, J., Lu, W., Wu, S., Cheng, Y., and Gouaux, E. (2015). Glycine receptor mechanism elucidated by electron cryo-microscopy. Nature 526, 224–229.

Durisic, N., Godin, A.G., Wever, C.M., Heyes, C.D., Lakadamyali, M., and Dent, J.A. (2012). Stoichiometry of the human glycine receptor revealed by direct subunit counting. The Journal of neuroscience : the official journal of the Society for Neuroscience 32, 12915–12920.

Eccles, J.C. (1982). The synapse: from electrical to chemical transmission. Annual review of neuroscience 5, 325–339.

Emsley, P., Lohkamp, B., Scott, W.G., and Cowtan, K. (2010). Features and development of Coot. Acta Crystallogr D Biol Crystallogr 66, 486–501.

Flayhan, A., Mertens, H.D.T., Ural-Blimke, Y., Martinez Molledo, M., Svergun, D.I., and Low, C. (2018). Saposin Lipid Nanoparticles: A Highly Versatile and Modular Tool for Membrane Protein Research. Structure 26, 345–355 e345.

Frauenfeld, J., Loving, R., Armache, J.P., Sonnen, A.F., Guettou, F., Moberg, P., Zhu, L., Jegerschold, C., Flayhan, A., Briggs, J.A., et al. (2016). A saposin-lipoprotein nanoparticle system for membrane proteins. Nat Methods 13, 345–351.

Fujii, Y., Kaneko, M., Neyazaki, M., Nogi, T., Kato, Y., and Takagi, J. (2014). PA tag: a versatile protein tagging system using a super high affinity antibody against a dodecapeptide derived from human podoplanin. Protein Expr Purif 95, 240–247.

Goehring, A., Lee, C.H., Wang, K.H., Michel, J.C., Claxton, D.P., Baconguis, I., Althoff, T., Fischer, S., Garcia, K.C., and Gouaux, E. (2014). Screening and large-scale expression of membrane proteins in mammalian cells for structural studies. Nat Protoc 9, 2574–2585.

Griffon, N., Buttner, C., Nicke, A., Kuhse, J., Schmalzing, G., and Betz, H. (1999). Molecular determinants of glycine receptor subunit assembly. The EMBO journal 18, 4711–4721.

Grudzinska, J., Schemm, R., Haeger, S., Nicke, A., Schmalzing, G., Betz, H., and Laube, B. (2005). The beta subunit determines the ligand binding properties of synaptic glycine receptors. Neuron 45, 727–739.

Harvey, R.J., Depner, U.B., Wassle, H., Ahmadi, S., Heindl, C., Reinold, H., Smart, T.G., Harvey, K., Schutz, B., Abo-Salem, O.M., et al. (2004). GlyR alpha3: an essential target for spinal PGE2-mediated inflammatory pain sensitization. Science 304, 884–887.

Huang, X., Chen, H., Michelsen, K., Schneider, S., and Shaffer, P.L. (2015). Crystal structure of human glycine receptor-alpha3 bound to antagonist strychnine. Nature 526, 277–280.

Kirsch, J., Wolters, I., Triller, A., and Betz, H. (1993). Gephyrin antisense oligonucleotides prevent glycine receptor clustering in spinal neurons. Nature 366, 745–748.

Kucukelbir, A., Sigworth, F.J., and Tagare, H.D. (2014). Quantifying the local resolution of cryo-EM density maps. Nat Methods 11, 63–65.

Kuhse, J., Laube, B., Magalei, D., and Betz, H. (1993). Assembly of the inhibitory glycine receptor: identification of amino acid sequence motifs governing subunit stoichiometry. Neuron 11, 1049–1056.

Kumar, A., Basak, S., Rao, S., Gicheru, Y., Mayer, M.L., Sansom, M.S.P., and Chakrapani, S. (2020). Mechanisms of activation and desensitization of full-length glycine receptor in lipid nanodiscs. Nature communications 11, 3752.

Langosch, D., Thomas, L., and Betz, H. (1988). Conserved quaternary structure of ligand-gated ion channels: the postsynaptic glycine receptor is a pentamer. Proceedings of the National Academy of Sciences of the United States of America 85, 7394–7398.

Li, L., Zhang, W., Liu, Q., Gao, Y., Gao, Y., Wang, Y., Wang, D.Z., Li, Z., and Wang, T. (2013). Structural Insights on the bacteriolytic and self-protection mechanism of muramidase effector Tse3 in Pseudomonas aeruginosa. J Biol Chem 288, 30607–30613.

Liebschner, D., Afonine, P.V., Baker, M.L., Bunkoczi, G., Chen, V.B., Croll, T.I., Hintze, B., Hung, L.W., Jain, S., McCoy, A.J., et al. (2019). Macromolecular structure determination using X-rays, neutrons and electrons: recent developments in Phenix. Acta Crystallogr D Struct Biol 75, 861–877.

Lynch, J.W. (2004). Molecular structure and function of the glycine receptor chloride channel. Physiological reviews 84, 1051–1095.

Lynch, J.W. (2009). Native glycine receptor subtypes and their physiological roles. Neuropharmacology 56, 303–309.

Lynch, J.W., and Callister, R.J. (2006). Glycine receptors: a new therapeutic target in pain pathways. Current opinion in investigational drugs 7, 48–53.

Malosio, M.L., Marqueze-Pouey, B., Kuhse, J., and Betz, H. (1991). Widespread expression of glycine receptor subunit mRNAs in the adult and developing rat brain. The EMBO journal 10, 2401–2409.

McGuire, H., Aurousseau, M.R., Bowie, D., and Blunck, R. (2012). Automating single subunit counting of membrane proteins in mammalian cells. The Journal of biological chemistry 287, 35912–35921.

Meyer, G., Kirsch, J., Betz, H., and Langosch, D. (1995). Identification of a gephyrin binding motif on the glycine receptor beta subunit. Neuron 15, 563–572.

Mohammadi, B., Krampfl, K., Cetinkaya, C., Moschref, H., Grosskreutz, J., Dengler, R., and Bufler, J. (2003). Kinetic analysis of recombinant mammalian alpha(1) and alpha(1)beta glycine receptor channels. Eur Biophys J 32, 529–536.

Morales-Perez, C.L., Noviello, C.M., and Hibbs, R.E. (2016). Manipulation of Subunit Stoichiometry in Heteromeric Membrane Proteins. Structure 24, 797–805.

Moss, S.J., and Smart, T.G. (2001). Constructing inhibitory synapses. Nature reviews Neuroscience 2, 240–250.

Mueller, P., Rudin, D.O., Tien, H.T., and Wescott, W.C. (1962). Reconstitution of cell membrane structure in vitro and its transformation into an excitable system. Nature 194, 979–980.

Patrizio, A., Renner, M., Pizzarelli, R., Triller, A., and Specht, C.G. (2017). Alpha subunit-dependent glycine receptor clustering and regulation of synaptic receptor numbers. Scientific reports 7, 10899.

Patrizio, A., and Specht, C.G. (2016). Counting numbers of synaptic proteins: absolute quantification and single molecule imaging techniques. Neurophotonics 3, 041805.

Pettersen, E.F., Goddard, T.D., Huang, C.C., Couch, G.S., Greenblatt, D.M., Meng, E.C., and Ferrin, T.E. (2004). UCSF Chimera--a visualization system for exploratory research and analysis. J Comput Chem 25, 1605–1612.

Pfeiffer, F., and Betz, H. (1981). Solubilization of the glycine receptor from rat spinal cord. Brain Res 226, 273–279.

Phulera, S., Zhu, H., Yu, J., Claxton, D.P., Yoder, N., Yoshioka, C., and Gouaux, E. (2018). Cryo-EM structure of the benzodiazepine-sensitive alpha1beta1gamma2S tri-heteromeric GABAA receptor in complex with GABA. eLife 7.

Pribilla, I., Takagi, T., Langosch, D., Bormann, J., and Betz, H. (1992). The atypical M2 segment of the beta subunit confers picrotoxinin resistance to inhibitory glycine receptor channels. The EMBO journal 11, 4305–4311.

Rahman, M.M., Teng, J., Worrell, B.T., Noviello, C.M., Lee, M., Karlin, A., Stowell, M.H.B., and Hibbs, R.E. (2020). Structure of the Native Muscle-type Nicotinic Receptor and Inhibition by Snake Venom Toxins. Neuron 106, 952–962 e955.

Sander, B., Tria, G., Shkumatov, A.V., Kim, E.Y., Grossmann, J.G., Tessmer, I., Svergun, D.I., and Schindelin, H. (2013). Structural characterization of gephyrin by AFM and SAXS reveals a mixture of compact and extended states. Acta crystallographica Section D, Biological crystallography 69, 2050–2060.

Schaefer, N., Roemer, V., Janzen, D., and Villmann, C. (2018). Impaired Glycine Receptor Trafficking in Neurological Diseases. Frontiers in molecular neuroscience 11, 291.

Scheres, S.H.W. (2012). RELION: Implementation of a Bayesian approach to cryo-EM structure determination. J Struct Biol 180, 519–530.

Schmidt, T., Schutz, G.J., Gruber, H.J., and Schindler, H. (1996). Local stoichiometries determined by counting individual molecules. Analytical chemistry 68, 4397–4401.

Schmieden, V., Grenningloh, G., Schofield, P.R., and Betz, H. (1989). Functional expression in Xenopus oocytes of the strychnine binding 48 kd subunit of the glycine receptor. The EMBO journal 8, 695–700.

Schneider, C.A., Rasband, W.S., and Eliceiri, K.W. (2012). NIH Image to ImageJ: 25 years of image analysis. Nat Methods 9, 671–675.

Schrödinger, L. (2015). The PyMOL molecular graphics system, version 2.3.1.

Shan, Q., Haddrill, J.L., and Lynch, J.W. (2001). A single beta subunit M2 domain residue controls the picrotoxin sensitivity of alphabeta heteromeric glycine receptor chloride channels. J Neurochem 76, 1109–1120.

Smart, O.S., Neduvelil, J.G., Wang, X., Wallace, B.A., and Sansom, M.S. (1996). HOLE: a program for the analysis of the pore dimensions of ion channel structural models. J Mol Graph 14, 354–360, 376.

Sola, M., Bavro, V.N., Timmins, J., Franz, T., Ricard-Blum, S., Schoehn, G., Ruigrok, R.W., Paarmann, I., Saiyed, T., O’Sullivan, G.A., et al. (2004). Structural basis of dynamic glycine receptor clustering by gephyrin. The EMBO journal 23, 2510–2519.

Sontheimer, H., Becker, C.M., Pritchett, D.B., Schofield, P.R., Grenningloh, G., Kettenmann, H., Betz, H., and Seeburg, P.H. (1989). Functional chloride channels by mammalian cell expression of rat glycine receptor subunit. Neuron 2, 1491–1497.

Specht, C.G., Izeddin, I., Rodriguez, P.C., El Beheiry, M., Rostaing, P., Darzacq, X., Dahan, M., and Triller, A. (2013). Quantitative nanoscopy of inhibitory synapses: counting gephyrin molecules and receptor binding sites. Neuron 79, 308–321.

Takahashi, T., Momiyama, A., Hirai, K., Hishinuma, F., and Akagi, H. (1992). Functional correlation of fetal and adult forms of glycine receptors with developmental changes in inhibitory synaptic receptor channels. Neuron 9, 1155–1161.

Usukura, E., Narita, A., Yagi, A., Ito, S., and Usukura, J. (2016). An Unroofing Method to Observe the Cytoskeleton Directly at Molecular Resolution Using Atomic Force Microscopy. Sci Rep 6, 27472.

Walker, P.A., Leong, L.E., Ng, P.W., Tan, S.H., Waller, S., Murphy, D., and Porter, A.G. (1994). Efficient and rapid affinity purification of proteins using recombinant fusion proteases. Biotechnology (N Y) 12, 601–605.

Wang, W., Touhara, K.K., Weir, K., Bean, B.P., and MacKinnon, R. (2016). Cooperative regulation by G proteins and Na(+) of neuronal GIRK2 K(+) channels. Elife 5.

Wang, W., Whorton, M.R., and MacKinnon, R. (2014). Quantitative analysis of mammalian GIRK2 channel regulation by G proteins, the signaling lipid PIP2 and Na+ in a reconstituted system. eLife 3, e03671.

Wassle, H., Heinze, L., Ivanova, E., Majumdar, S., Weiss, J., Harvey, R.J., and Haverkamp, S. (2009). Glycinergic transmission in the Mammalian retina. Frontiers in molecular neuroscience 2, 6.

Weltzien, F., Puller, C., O’Sullivan, G.A., Paarmann, I., and Betz, H. (2012). Distribution of the glycine receptor beta-subunit in the mouse CNS as revealed by a novel monoclonal antibody. The Journal of comparative neurology 520, 3962–3981.

Werman, R., Davidoff, R.A., and Aprison, M.H. (1967). Inhibition of motoneurones by iontophoresis of glycine. Nature 214, 681–683.

Williams, C.J., Headd, J.J., Moriarty, N.W., Prisant, M.G., Videau, L.L., Deis, L.N., Verma, V., Keedy, D.A., Hintze, B.J., Chen, V.B., et al. (2018). MolProbity: More and better reference data for improved all-atom structure validation. Protein Sci 27, 293–315.

Yang, Z., Cromer, B.A., Harvey, R.J., Parker, M.W., and Lynch, J.W. (2007). A proposed structural basis for picrotoxinin and picrotin binding in the glycine receptor pore. J Neurochem 103, 580–589.

Yang, Z., Taran, E., Webb, T.I., and Lynch, J.W. (2012). Stoichiometry and subunit arrangement of alpha1beta glycine receptors as determined by atomic force microscopy. Biochemistry 51, 5229–5231.

Yu, J., Zhu, H., Lape, R., Greiner, T., Du, J., Lu, W., Sivilotti, L., and Gouaux, E. (2021). Mechanism of gating and partial agonist action in the glycine receptor. Cell.

Yu, T.W., Chahrour, M.H., Coulter, M.E., Jiralerspong, S., Okamura-Ikeda, K., Ataman, B., Schmitz-Abe, K., Harmin, D.A., Adli, M., Malik, A.N., et al. (2013). Using whole-exome sequencing to identify inherited causes of autism. Neuron 77, 259–273.

Zacchi, P., Antonelli, R., and Cherubini, E. (2014). Gephyrin phosphorylation in the functional organization and plasticity of GABAergic synapses. Frontiers in cellular neuroscience 8, 103.

Zacharias, D.A., Violin, J.D., Newton, A.C., and Tsien, R.Y. (2002). Partitioning of lipid-modified monomeric GFPs into membrane microdomains of live cells. Science 296, 913–916.

Zeng, M., Chen, X., Guan, D., Xu, J., Wu, H., Tong, P., and Zhang, M. (2018). Reconstituted Postsynaptic Density as a Molecular Platform for Understanding Synapse Formation and Plasticity. Cell 174, 1172–1187 e1116.

Zhang, K. (2016). Gctf: Real-time CTF determination and correction. J Struct Biol 193, 1–12.

Zheng, S.Q., Palovcak, E., Armache, J.P., Verba, K.A., Cheng, Y.F., and Agard, D.A. (2017). MotionCor2: anisotropic correction of beam-induced motion for improved cryo-electron microscopy. Nature Methods 14, 331–332.

Zhu, S., Noviello, C.M., Teng, J., Walsh, R.M., Jr., Kim, J.J., and Hibbs, R.E. (2018). Structure of a human synaptic GABAA receptor. Nature 559, 67–72.

